# An expanded phylogeny of *Setaria* (Poaceae, Panicoideae, Paniceae) and its relationships within the subtribe Cenchrinae

**DOI:** 10.1101/2022.08.22.504865

**Authors:** Carolina Delfini, Juan M. Acosta, José F. Pensiero, Sandra S. Aliscioni

## Abstract

*Setaria* P. Beauv. is the largest genus of the “bristle clade”, including between 115 and 160 species. Previous molecular phylogenetic studies showed *Setaria* likely to be para- or polyphyletic, retrieving several clades apparently consistent in all analyses and correlated with the geographic origin of species. In this study, we evaluate the phylogeny of the subtribe Cenchrinae using parsimony, likelihood, and Bayesian inference based on the plastid marker *ndh*F and increasing the number of sampled species. Our main objective was analyze American taxa with inflorescences of the “Paspalidium type” (i.e., subgenera *Paurochaetium* and *Reverchoniae*) to test whether they, as traditionally circumscribed, form a natural group. Our findings recovered both subgenera as polyphyletic, with their species distributed in different morphologically distinctive clades and not necessarily correlated with the geographic origin. Additionally, we were able to include a second voucher of species that were imprecisely located in previous studies and define their placements in the tree, as well as confirm that *Setaria* is polyphyletic as currently delineated. A comparison with the results from other studies, comments on *Stenotaphrum* Trin. and a brief discussion on conflicting placements in the “Cenchrus clade”, and of *Acritochaete* Pilg. are also included here.

## Introduction

*Setaria* P. Beauv. is a cosmopolitan genus of grasses comprising between 115 and 160 species [1–2], commonly found in open environments and woodlands [2–5]. The species grow mostly in tropical and subtropical latitudes, though several are present in cold regions of both hemispheres [2–5]. The Old World species are concentrated in tropical Africa, including 12 endemic to Madagascar [6–7], whereas in the New World the center of species diversity is Brazil [3] with 30 native [8–9].

*Setaria* is one of several genera in the subtribe Cenchrinae [10–12] that is characterized by having highly modified inflorescences with sterile branches, often known as setae or “bristles” persist when the spikelets fall at maturity [2, 5, 13]. Despite being a morphologically well-characterized genus, the phylogeny of *Setari*a and its related genera lacks resolution mainly in defining relationships among large clades [13–14]. The most complete phylogeny at the present [14] is based on the plastid marker *ndh*F and shows the genus likely to be para- or polyphyletic, with several moderately to strongly supported clades apparently consistent in all analyses [14]. These clades represent linages correlated basically to geographic distribution but, relationships among them are unclear. In addition, other small genera of Cenchrinae (e.g., *Ixophorus* Schltdl., *Setariopsis* Scribn. ex Millsp., *Spinifex* L., *Uranthoecium* Stapf, *Zygochloa* S.T. Blake, and *Zuloagaea* E. Bess) are consistently resolved within *Setaria*, making morphological affinities between the species even more difficult to establish [14]. Based on the preceding findings [14] plus new taxa added in [15], [13] presented a phylogeny of subtribe Cenchrinae, focusing on *Setaria* species. In this tree, four blocks including clades, groups of clades or ungrouped species are indicated and named by the geographic origin (i.e., 1. Africa, tropical-Asia, 2. Australia, Australasia, 3.temperate Asia, and 4. Americas). The lack of a well-resolved phylogeny along with the difficult morphological delimitation of some species mean that *Setaria* requires further in depth research. It is clearly not a natural group but more evidence is still needed to allow restructuring of the taxonomy of *Setaria* in association with the other genera within the subtribe Cenchrinae [13– 14].

Previous authors have had differing opinions regarding the infrageneric classification of *Setaria*, but in general, one or two distinctive groups of species have been recognized and the remainder placed in subgenus/section *Setaria* [3, 5, 16]. For tropical Africa species, [6] recognized four sections, that is, *Eu-setaria* Stapf characterized by its young blades not plicate and panicles usually spike-like, *Ptychophyllum* (A. Braun) Stapf including plants with plicate blades and open panicles, and sections *Panicatrix* Stapf & C.E. Hubb. and *Cymbosetaria* Stapf Hubb., distinguished by having rounded and keeled upper lemmas, respectively, and non-cylindrical inflorescences [6]. In addition to these four, [16] also recognized section *Setaria*, characterized by the blades not plicate and bristles usually below all the spikelets.

Using similar criteria and also based on the position of the bristles along the inflorescences, [3] recognized three subgenera for the North American species: *Setaria, Ptychophyllum* and *Paurochaetium* (Hitchc. & Chase) Rominger, which groups species with bristles present only at the ends of primary branches. [5], in his treatment of the South American species, recognized the subgenera *Setaria, Ptychophyllum* and proposed the new monotypic subgenus *Cernatum* Pensiero, to accommodate *Setaria cernua* Kunth, whose position in the infrageneric classification had long been uncertain.

In some species, the primary branches of the inflorescence are themselves unbranched (i.e., the spikelets are born directly on the primary branches) and these branches end in a bristle [2]. The Old World species with this type of inflorescences, called informally “Paspalidium type”, were placed in the genus *Paspalidium* Stapf. [17]. However, American species with similar type of inflorescences were treated in three different subgenera within *Setaria*: *Paurochaetium, Reverchoniae* W.E. Fox (segregated from subgenus *Paurochaetium*), and *Cernatum* [3, 5, 18].

Advances in morphological and systematic studies in *Setaria* have shown that there is another type of inflorescence in which some other spikelets (but not all) are associated with bristles in addition to the uppermost ones in the branch tip, although the general pattern is similar to the “Paspalidium type” [2, 13]. The recognition of this intermediate pattern led to the transfer of Old World *Paspalidium* species back to *Setaria* [1–2, 19–20], a result partially supported by molecular phylogenies [10–12, 14], and morphological and foliar anatomical data [21].

*Setaria* is currently recognized as a difficult and non-monophyletic genus; its species are isolated or segregate into many clades in the subtribe Cenchrinae and none of the tested subgenera (i.e., *Ptychophyllum* (A. Braun) Hitchc. and *Setaria*) are monophyletic. Besides the monotypic South American subgenus *Cernatum* analyzed by [14], whose placement conflicts in different analyses and is, up to now, unresolved, none American *Setaria* species with a “Paspalidium type” inflorescence (i.e., subgenera *Paurochaetium* and *Reverchoniae*) were sampled in existing phylogenies. Based on these previous results, our principal objective was include species of the subgenera *Paurochaetium* and *Reverchoniae* to test their positions, assuming a priori that these subgenera either would be resolved within the “American groups” due to their geographic origin, or related to species originally considered as *Paspalidium*, given their morphological similarities. Additionally, we added *ndh*F sequences of Old World species of *Setaria* and other genera of Cenchrinae from Genbank not considered in previous phylogenies [14]. We included new sequences of some other American species of *Setaria* (not *Paurochaetium* and *Reverchoniae*) and a new second voucher of some species that were imprecisely located in [14], as they were represented by partial, not fully double-stranded and/or poor-quality accessions, and their positions in the tree are now defined.

## Material and Methods

### Taxon sampling

The aligned data matrix used in the phylogenetic analyses includes a total of 178 accessions, of which 170 are ingroup corresponding to the subtribe Cenchrinae (Table 1). The chloroplast DNA (cpDNA) *ndh*F matrix previously published [14], excluding the outgroup, was augmented with 61 new sequences, of which 32 are *Setaria* (Table 1). Of these, we have sequenced 18 that corresponding to species of the subgenera *Paurochaetium* and *Reverchoniae* (Table 2) plus those without a defined placement in [14] (Table 1, indicated with **). Eight species belonging to six closely related genera were selected as outgroup, based on [10, 14]: *Aakia* J.R. Grande, *Eriochloa* Kunth, *Moorochloa* Veldkamp, *Panicum* L, *Rupichloa* Salariato & Morrone, and *Urochloa* P. Beauv. Information about vouchers and accession numbers of the new sequences obtained for this study and those available in GenBank are given in Table 1.

**Table 1.**
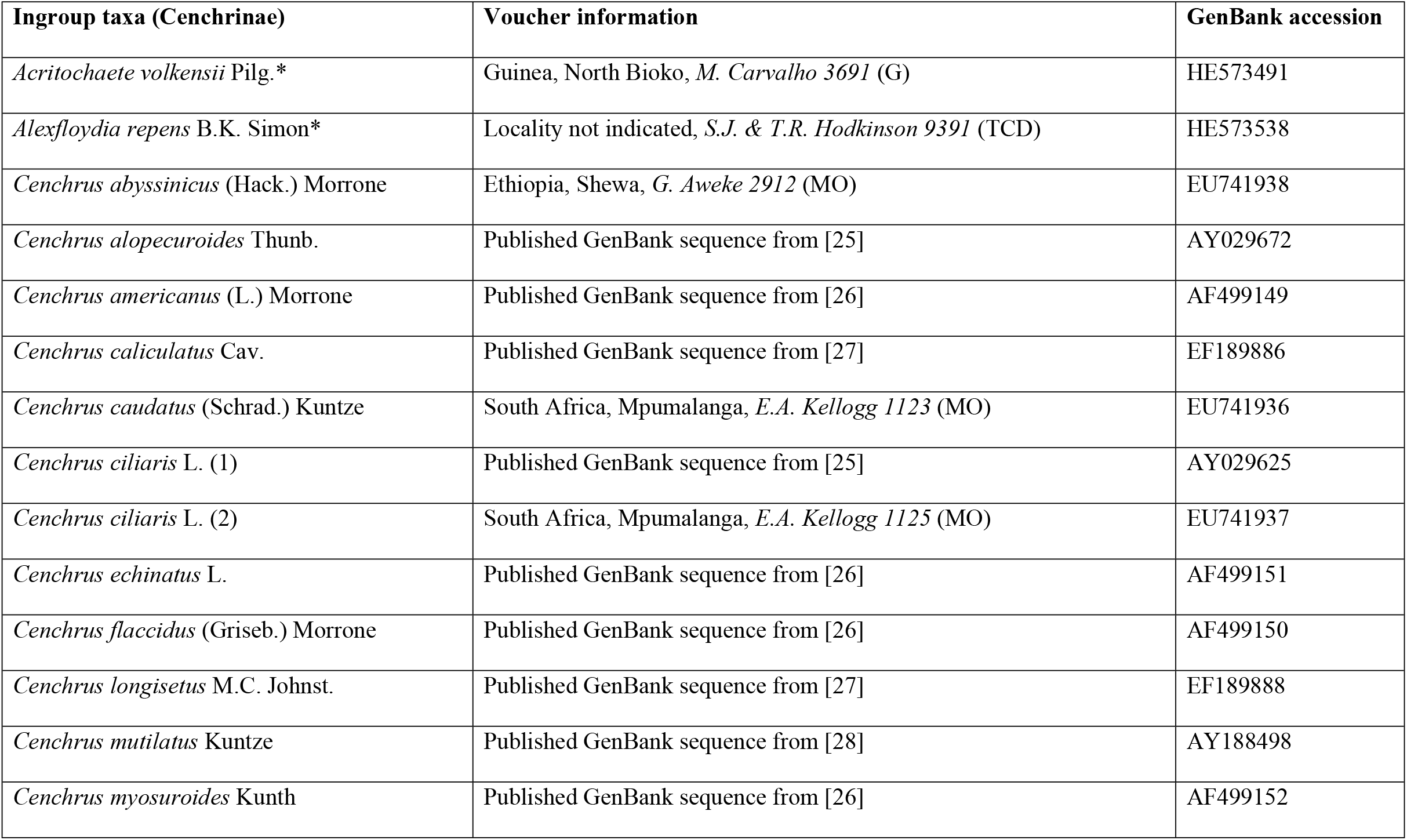
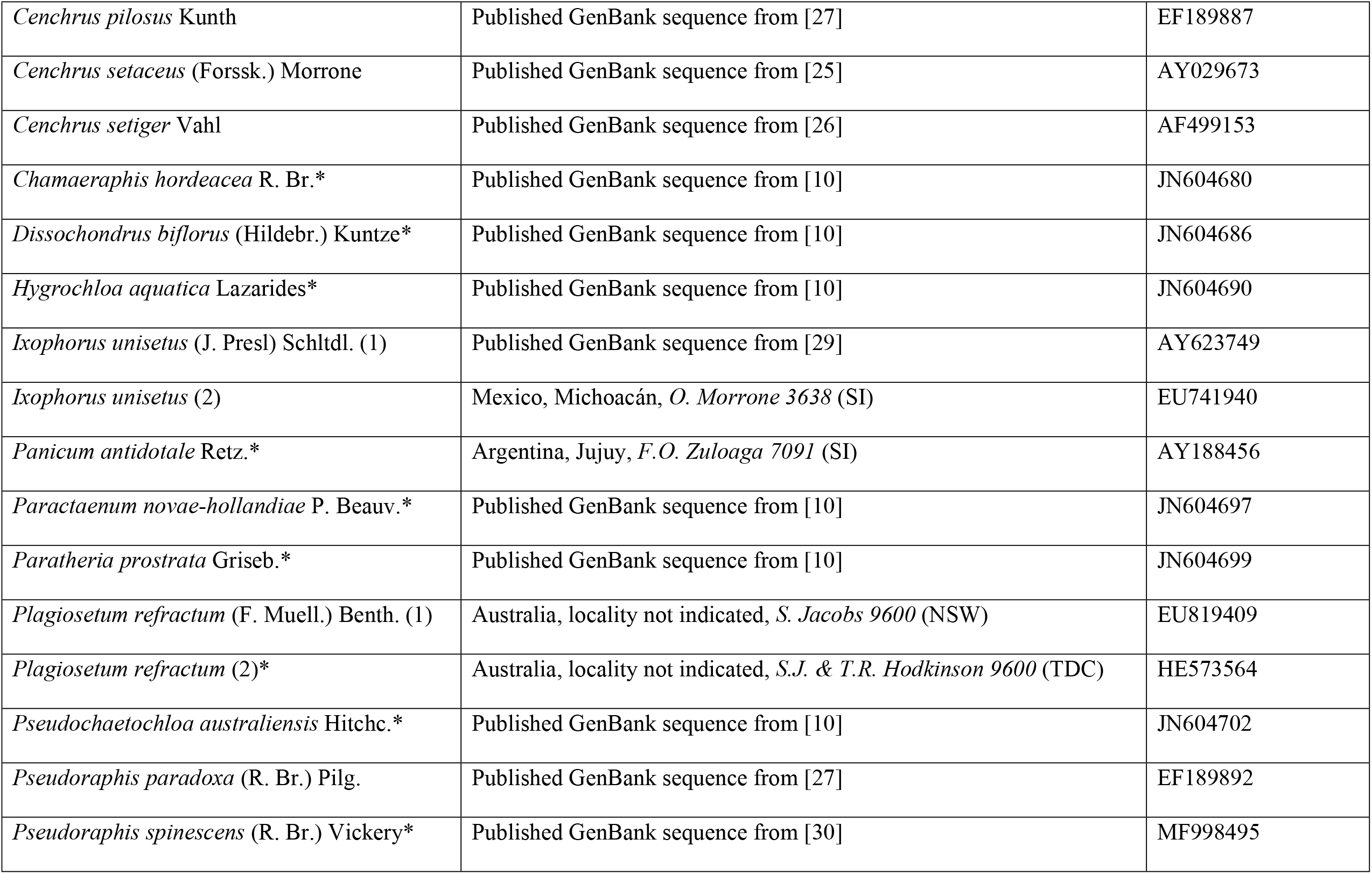
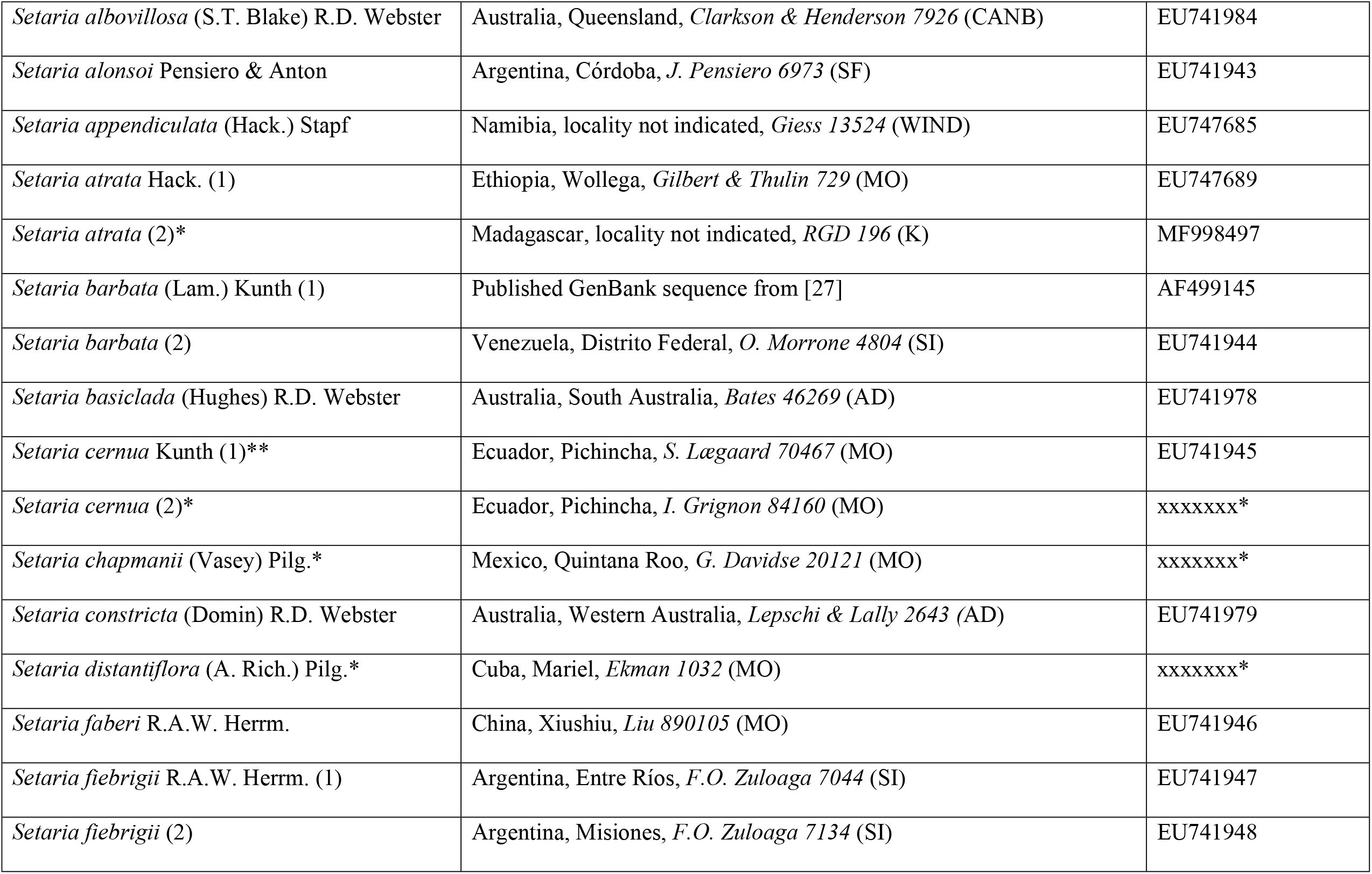
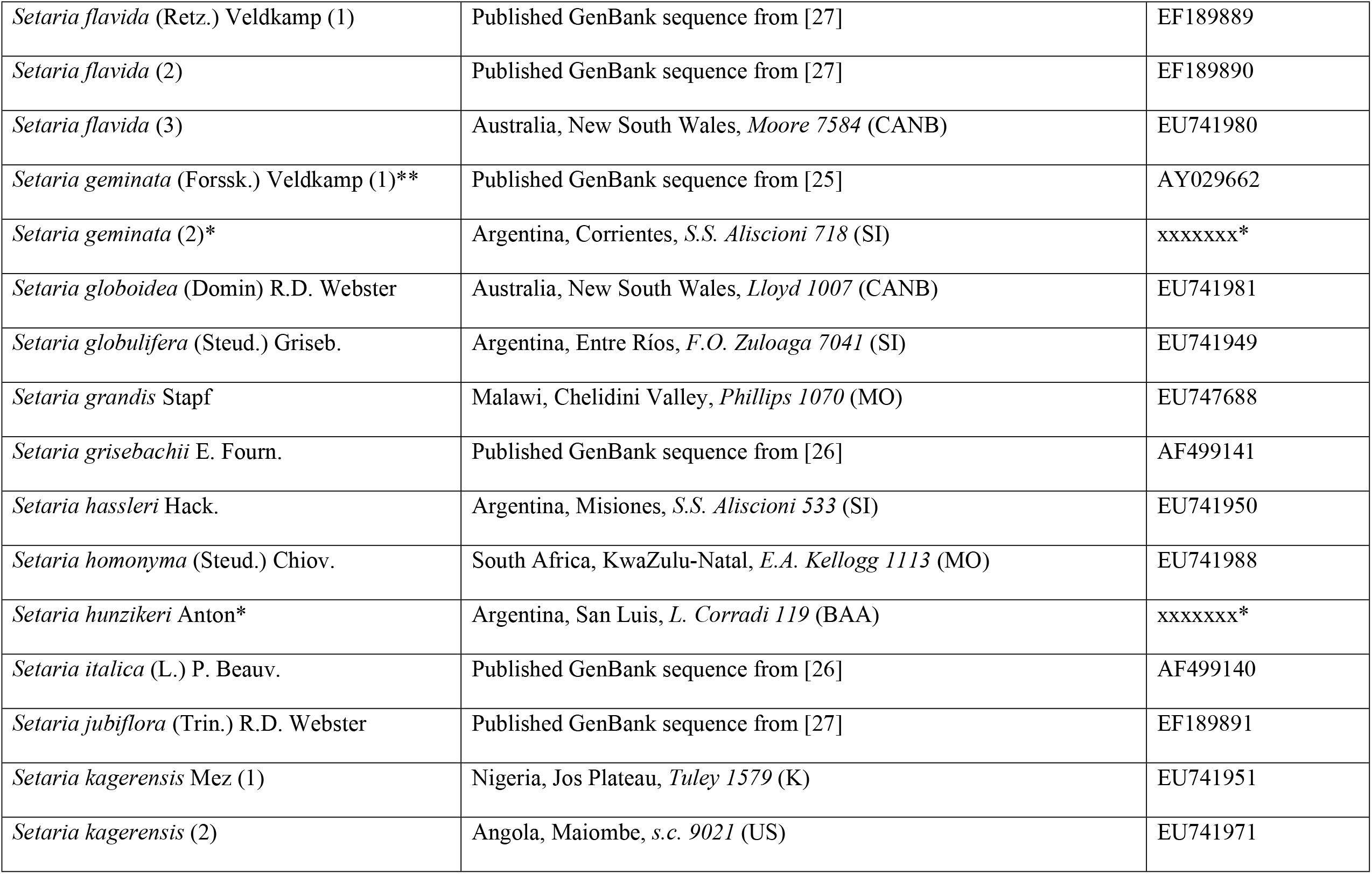
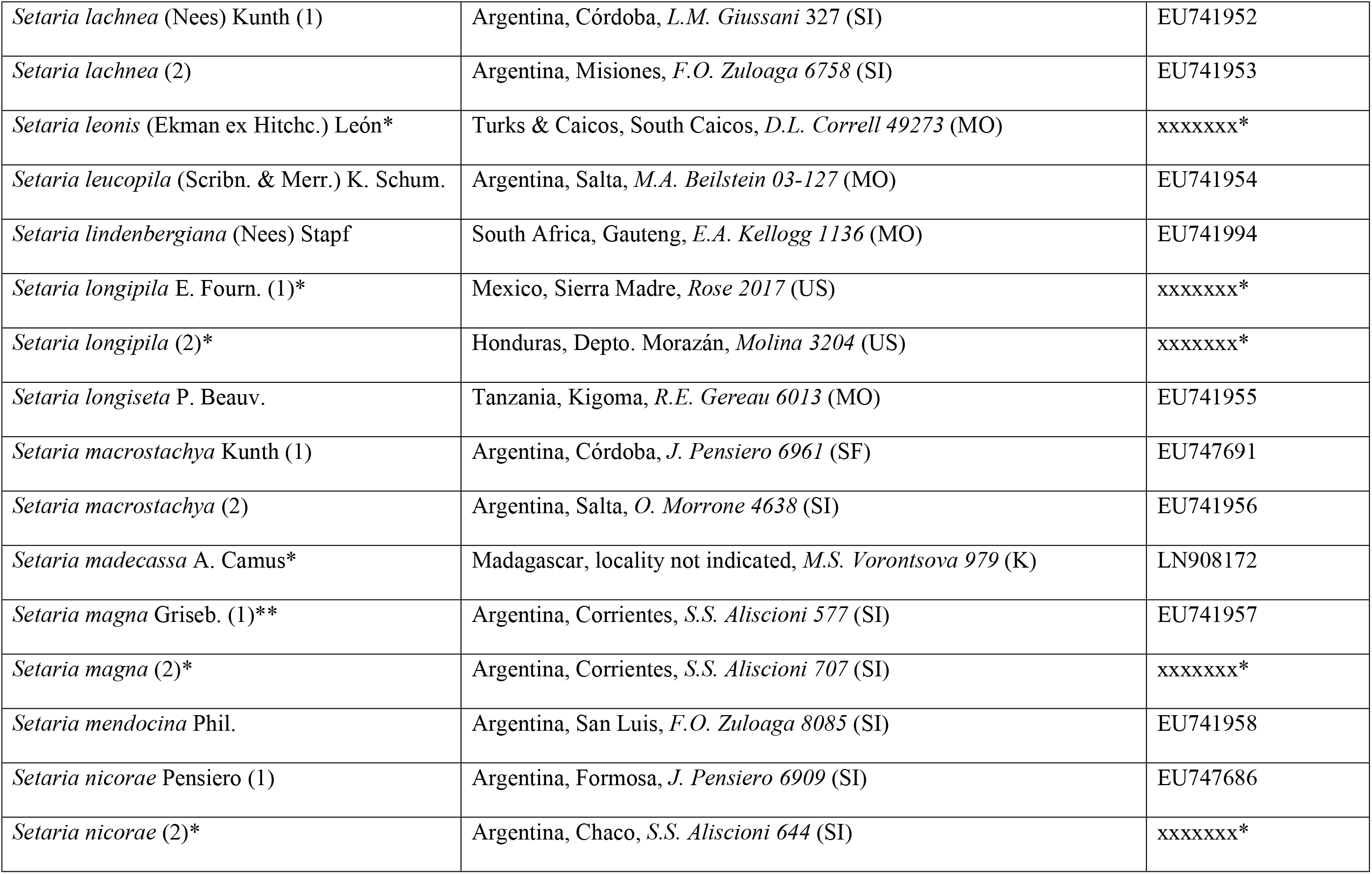
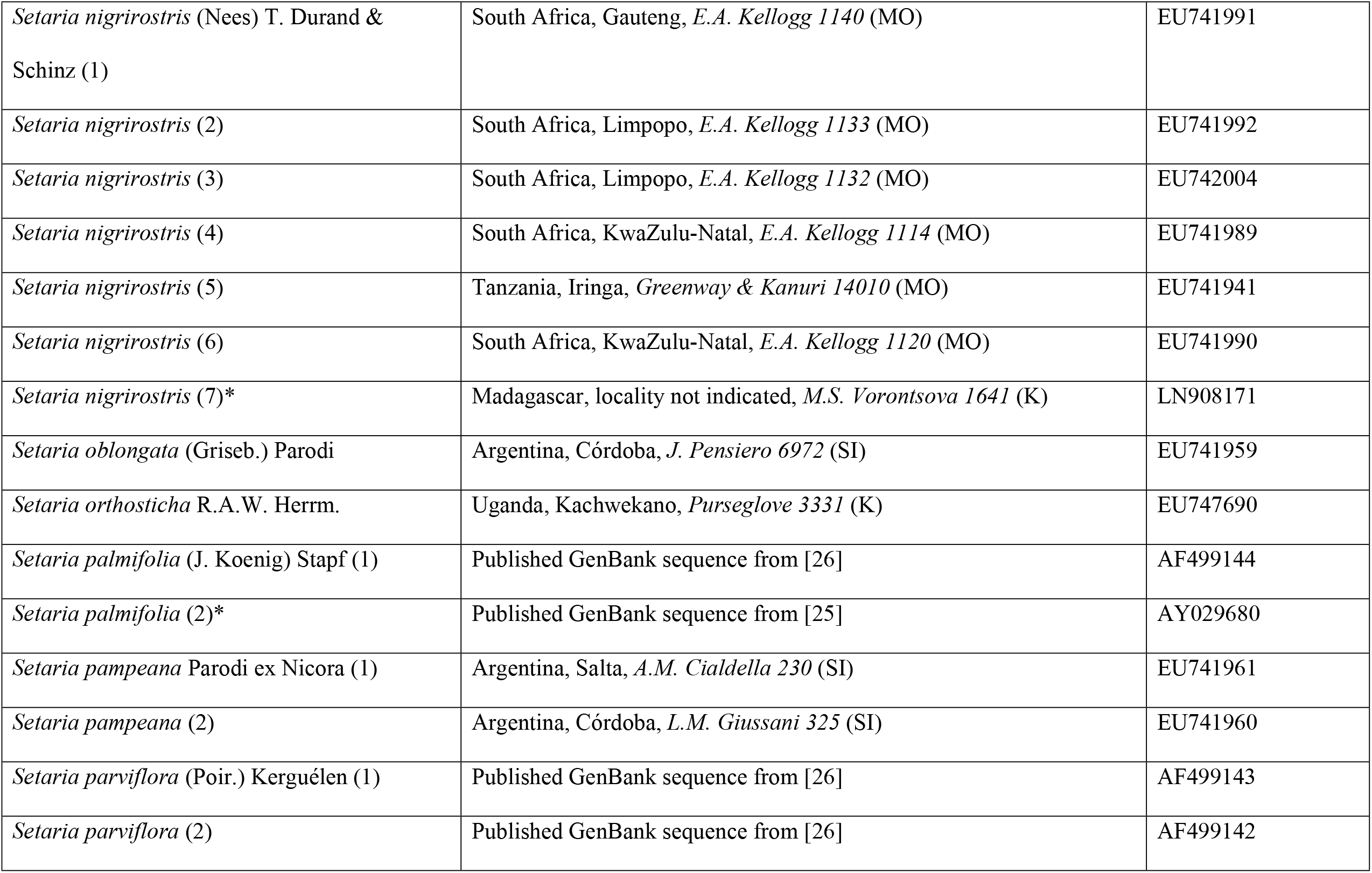
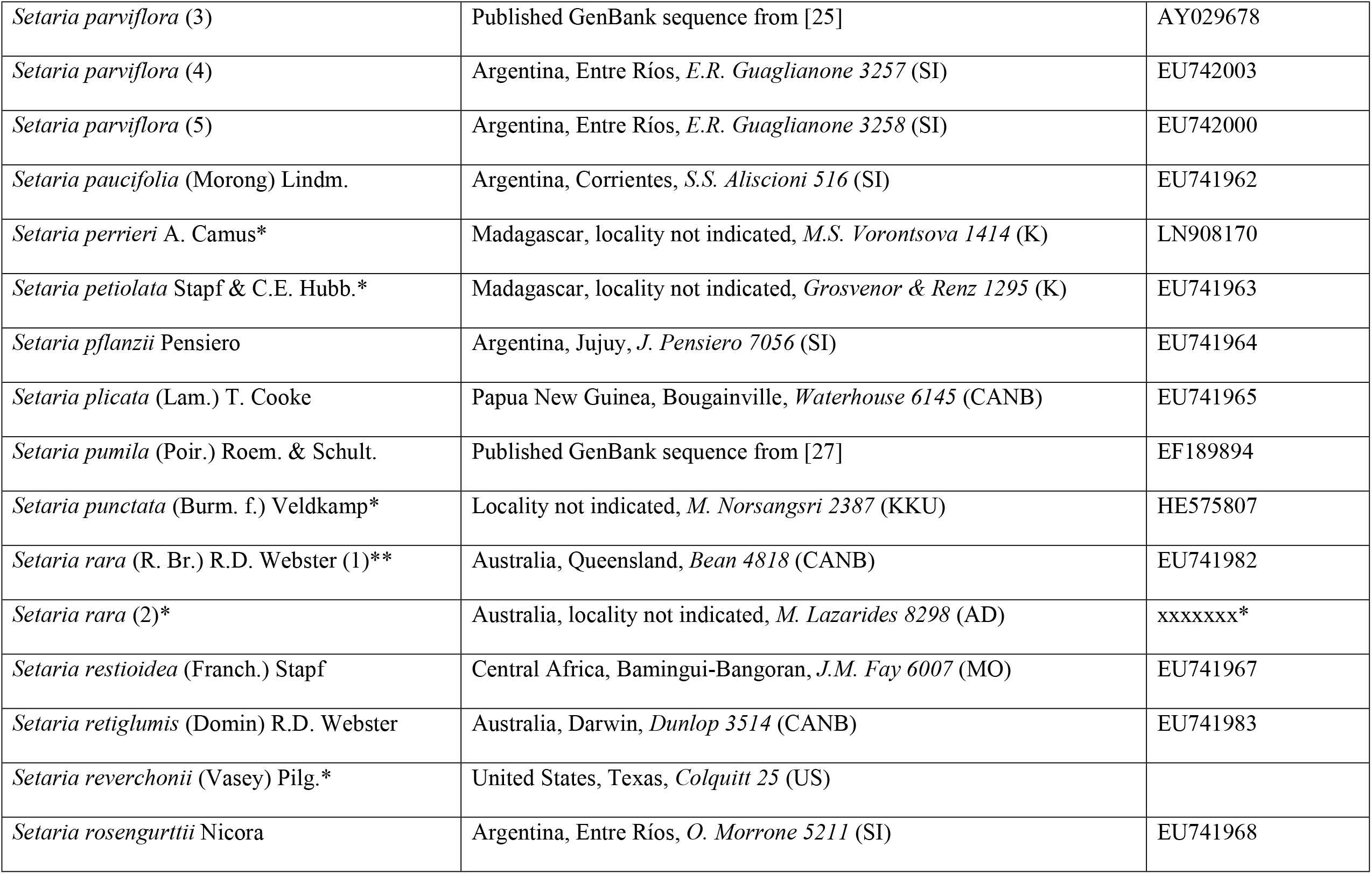
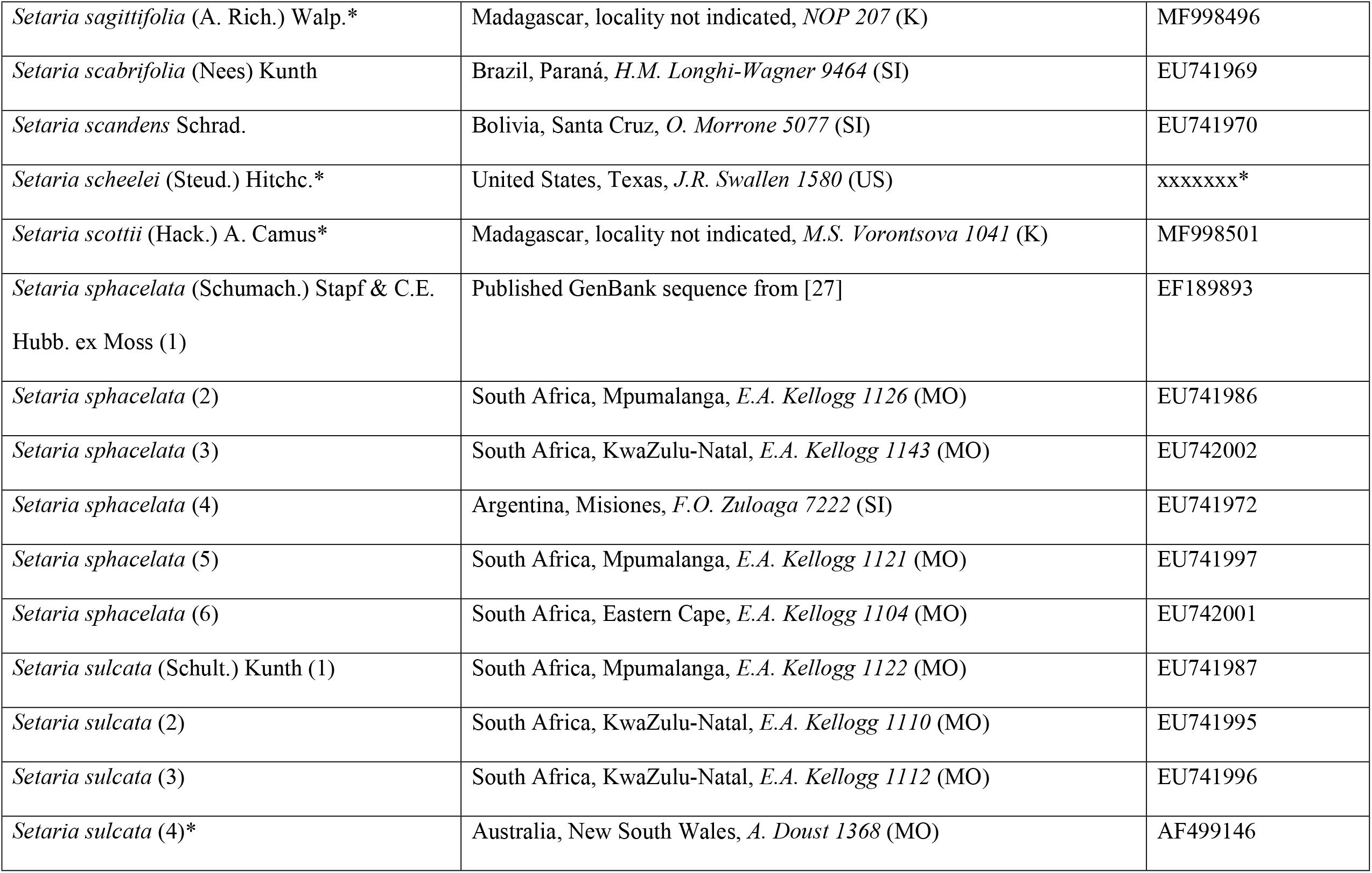
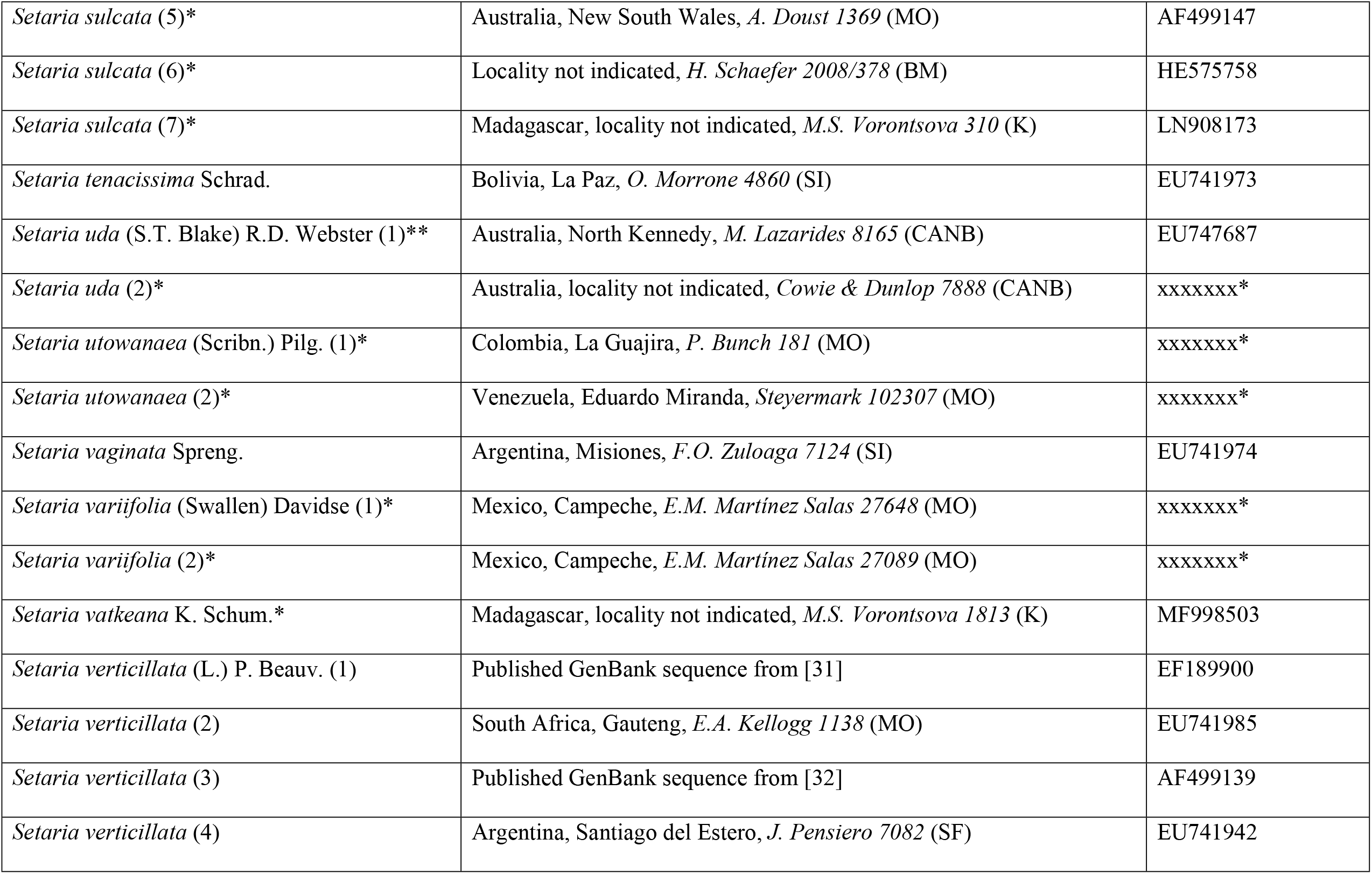
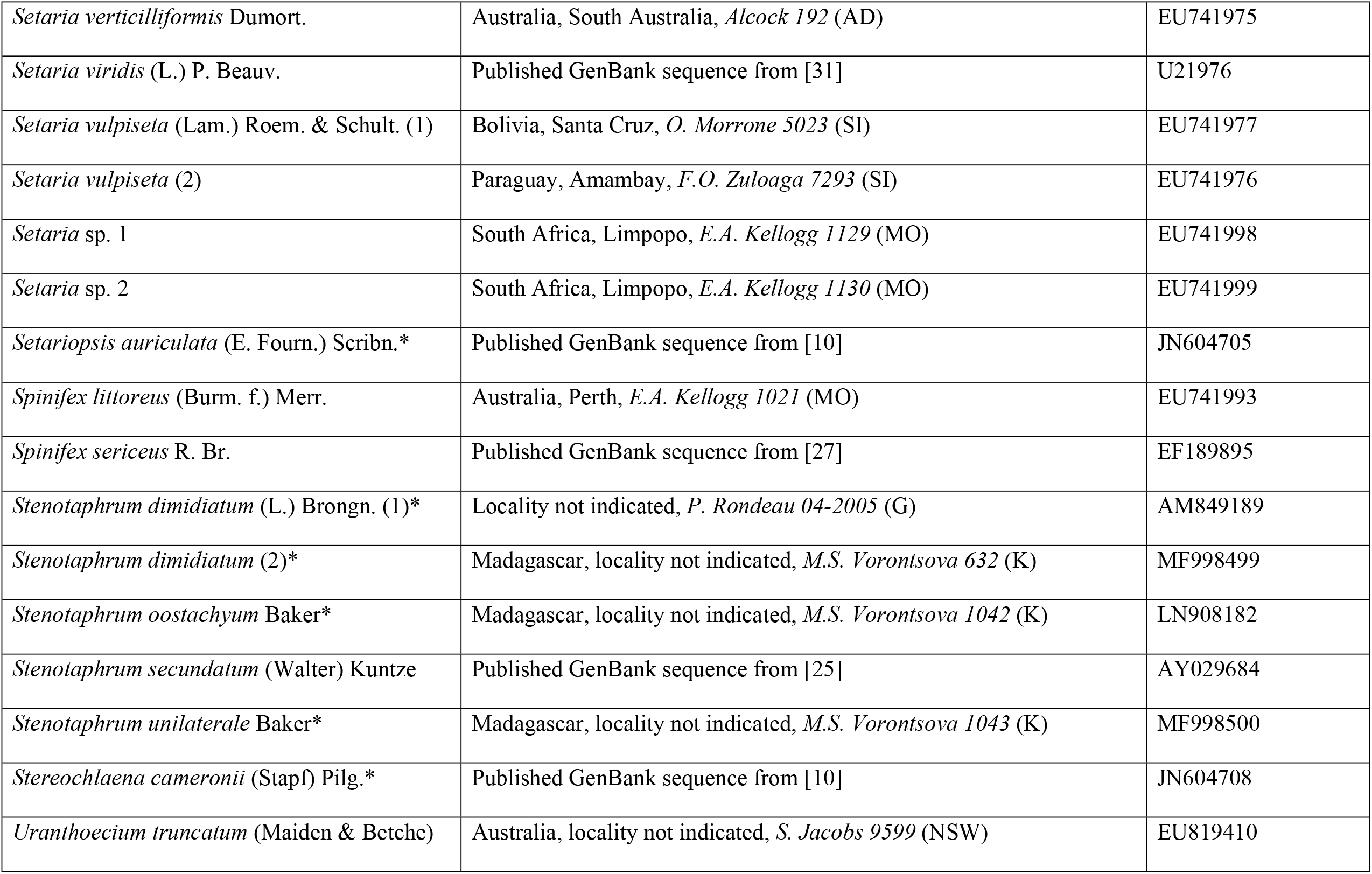
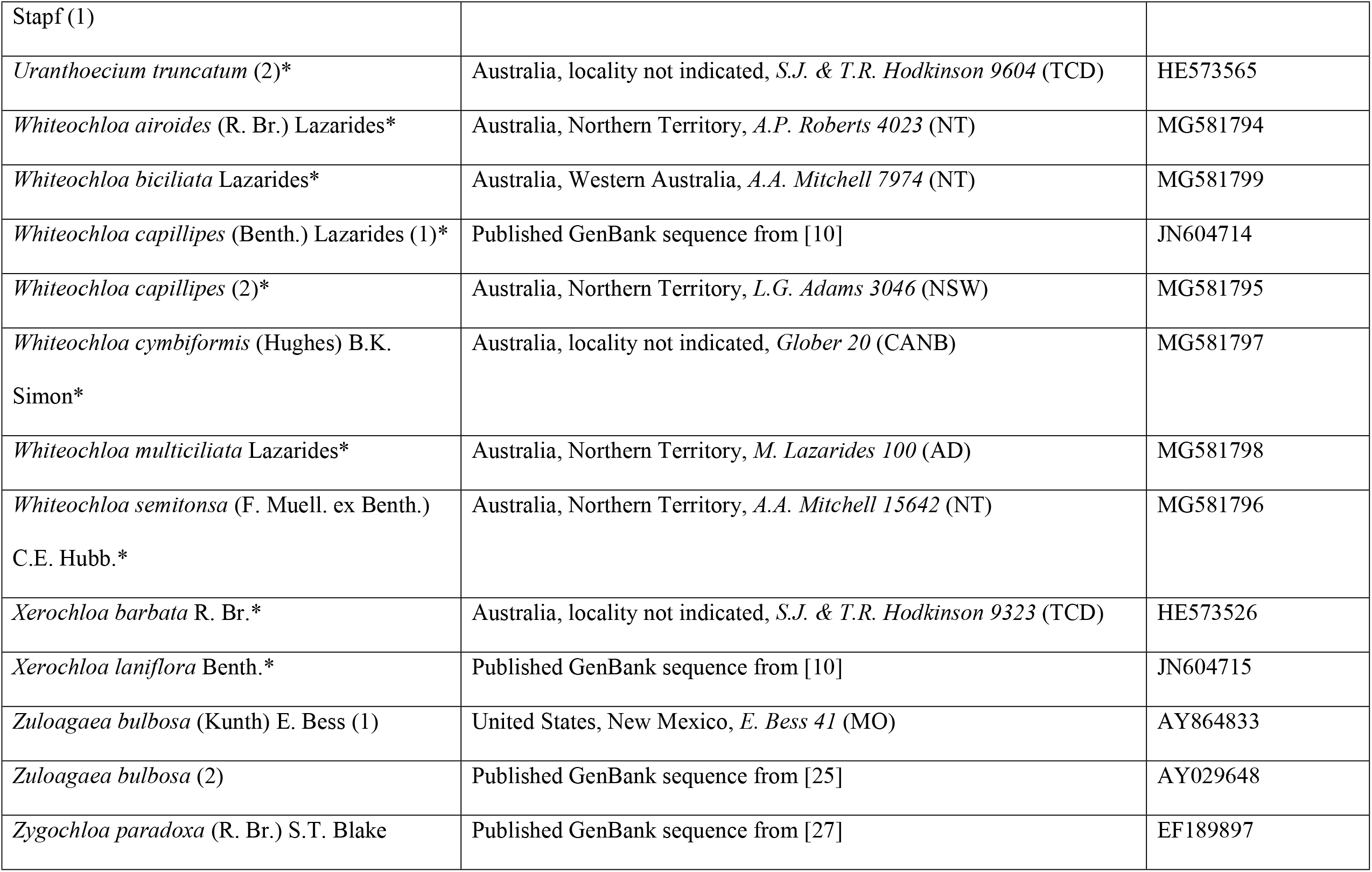
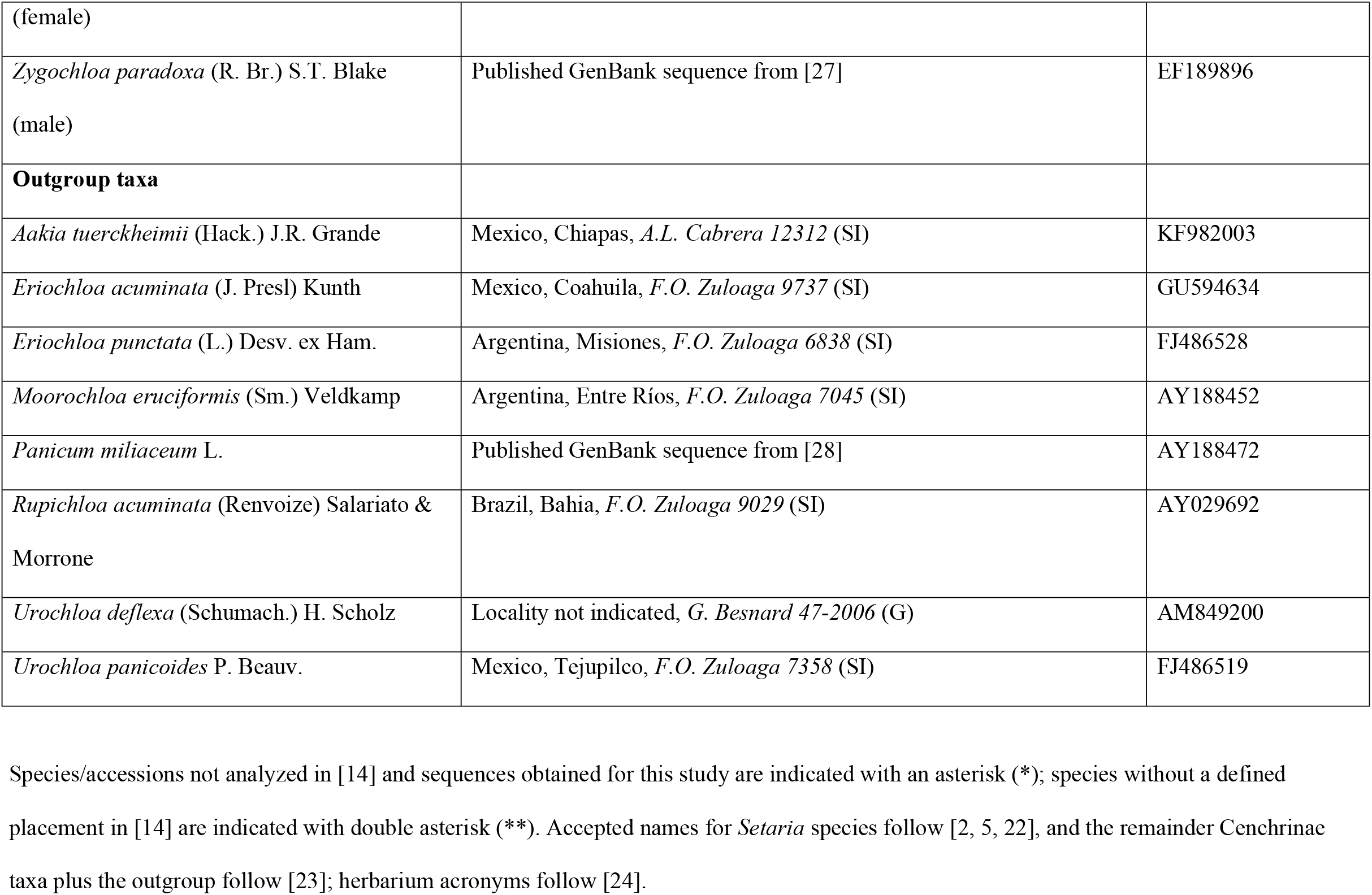
Taxa, voucher information, and GenBank accession numbers for *ndh*F sequences.

**Table 2.** Comparison of different classifications for the taxa of subgenera *Paurochaetium* and *Reverchoniae*, and their currently accepted names according to [22].

| [33] | [3] | [34] | [5] | [18] |  | Accepted names |
| --- | --- | --- | --- | --- | --- | --- |
| *Panicum* subg.<br>*Paurochaetium*<br>Hitchc. & Chase | *Setaria* subg.<br>*Paurochaetium*<br>(Hitchc. & Chase)<br>Rominger | *Paspalidium* Stapf | *Setaria* subg.<br>*Setaria* | *Setaria* subg.<br>*Paurochaetium*<br>(Hitchc. & Chase)<br>Rominger | *Setaria* subg.<br>*Reverchoninae*<br>W.E. Fox |  |
| *Panicum*<br>*chapmanii* Vasey | *Setaria chapmanii*<br>(Vasey) Pilg. | *Paspalidium*<br>*chapmanii* (Vasey)<br>R.W. Pohl | *Paspalidium*<br>*chapmanii* (Vasey)<br>R.W. Pohl | *Setaria chapmanii*<br>(Vasey) Pilg. |  | *Setaria chapmanii*<br>(Vasey) Pilg. |
| *Panicum*<br>*distantiflorum* A.<br>Rich. | *Setaria distantiflora*<br>(A. Rich.) Pilg. | *Paspalidium*<br>*distantiflorum* (A.<br>Rich.) Davidse &<br>R.W. Pohl | *Setaria distantiflora*<br>(A. Rich.) Pilg. | *Setaria*<br>*distantiflora* (A.<br>Rich.) Pilg. |  | *Setaria distantiflora*<br>(A. Rich.) Pilg. |
| *Panicum leonis*<br>Ekman ex Hitchc. | *Setaria leonis*<br>(Ekman ex Hitchc.)<br>León | *Paspalidium leonis*<br>(Ekman ex Hitchc.)<br>Davidse & R.W. Pohl | *Setaria leonis*<br>(Ekman ex Hitchc.)<br>León | *Setaria leonis*<br>(Ekman ex Hitchc.)<br>León |  | *Setaria leonis* (ex<br>Hitchc.) León |
| *Panicum pradatum* León ex Hitchc. | *Setaria pradana* (León ex Hitchc.) León | *Paspalidium pradatum* (León ex Hitchc.) Davidse & R.W. Pohl | *Setaria pradana* (León ex Hitchc.) León | *Setaria pradana* (León ex Hitchc.) León |  | *Setaria pradana* (León ex C.L. Hitchc.) León |
| *Panicum utowanaeum* Scribn. | *Setaria utowanaea* (Scribn.) Pilg. | *Paspalidium utowanaeum* (Scribn.) Davidse & R.W. Pohl | *Setaria utowanaea* (Scribn.) Pilg. | *Setaria utowanaea* var. *utowanaea* (Scribn.) Pilg. |  | *Setaria utowanaea* var. *utowanaea* (Scribn.) Pilg. |
| *Panicum ophiticola* Hitchc. & Ekman | *Setaria ophiticola* (Hitchc. & Ekman) León | *Paspalidium ophiticola* (Hitchc. & Ekman) Davidse & R.W. Pohl | *Setaria ophiticola* (Hitchc. & Ekman) León | *Setaria utowanaea* var. *ophiticola* (Hitchc. & Ekman) W.E. Fox |  | *Setaria utowanaea* var. *ophiticola* (Hitchc. & Ekman) W.E. Fox. |
|  | *Setaria subtransiens* Hitchc. & Ekman | *Paspalidium subtransiens* (Hitchc. & Ekman) Davidse & R.W. Pohl | *Setaria subtransiens* Hitchc. & Ekman | *Setaria utowanaea* var. *subtransiens* (Hitchc. & Ekman) W.E. Fox |  | *Setaria utowanaea* var. *subtransiens* (Hitchc. & Ekman) W.E. Fox |
| *Panicum reverchonii* Vasey | *Setaria reverchonii* (Vasey) Pilg. |  | *Setaria reverchonii* (Vasey) Pilg. |  | *Setaria reverchonii* subsp. | *Setaria reverchonii* subsp. *reverchonii* |
|  |  |  |  |  | *reverchonii*<br>Vasey) Pilg. | (Vasey) Pilg. |
| *Panicum firmulum*<br>Hitchc. & Chase | *Setaria firmula*<br>(Hitchc. & Chase)<br>Pilg. |  | *Setaria firmula*<br>(Hitchc. & Chase)<br>Pilg. |  | *Setaria*<br>*reverchonii* subsp.<br>*firmula* (Hitchc. &<br>Chase) W.E. Fox | *Setaria reverchonii*<br>subsp. *firmula*<br>(Hitchc. & Chase)<br>W.E. Fox |
| *Panicum*<br>*ramisetum* Scribn. | *Setaria ramiseta*<br>(Scribn.) Pilg. |  | *Setaria ramiseta*<br>(Scribn.) Pilg. |  | *Setaria*<br>*reverchonii* subsp.<br>*ramiseta* (Scribn.)<br>W.E. Fox | *Setaria reverchonii*<br>subsp. *ramiseta*<br>(Scribn.) W.E. Fox |
|  |  |  |  |  | *Setaria variifolia*<br>(Swallen) Davidse | *Setaria variifolia*<br>(Swallen) Davidse |

### DNA amplification and sequencing

Total genomic DNA was extracted from herbarium material using modified CTAB protocols from [35]. For the species that failed this protocol, the DNA was isolated using the DNeasy Plant Mini Kit (Qiagen, Hilden, Germany), following the manufacturer’s recommendations. Each species was amplified from a single voucher specimen but, a second voucher was also included for some taxa. The *ndh*F gene, coding NADH dehydrogenase subunit F, was amplified by polymerase chain reaction (PCR) and sequenced for each taxon. The complete region was amplified with a battery of primers in different combinations in four overlapping fragments using primer pairs specified by [28, 36]: 5F–536R, 536F–972R, 972F–1666R, and 1666F–3R. Due to a lot of samples with a difficult amplification of the region 1666F–3R, a new reverse primer near the 3R region was designed for PCR amplification and sequencing of *ndh*F within the subfamily Panicoideae: 2150R (5’– TCTCCKATACAAAAACYARCAAKAC–3’).

PCR reactions were performed in a 25 µl final volume with 50–100 ng of template DNA, 5 µl Green Promega GoTaq^®^ buffer (5 u/µl), 0.5 µl MgCl_2_ (25 mM), 1.25 µl dNTP (10 mM), 1 µl of each primer (10 pM), and 0.3 µl of Taq polymerase (5 u/µl) provided by Promega (Madison, Wisconsin, U.S.A.). Variations in MgCl_2_ (0.5–1 µl) and total DNA dilutions (1:5, 1:10 and 1:50) were used. The reactions were carried out using the following parameters: one cycle of 95 °C for 2 min, 39 cycles of 95 °C for 30 s, 48 °C for 30 s, and 72 °C for 1.5 min, and a final extension cycle of 72 °C for 10 min. A negative control with no template was included for each series of amplifications to eliminate the possibility of contamination. PCR products were run out on a 1% TBE (Tris-Borate-EDTA) agarose gel stained with SYBR Safe DNA gel stain (Invitrogen Life Technologies) and visualized in a blue-light transilluminator. PCR products were purified and automated sequencing was performed by Macrogen, Inc. (Seoul, South Korea). Forward and reverse strands were sequenced for all fragments, with a minimum overlap of 80%.

### Phylogenetic analyses

Sequence editing and assembly were performed with MEGA v. 7.0 [37]. Accuracy of sequences was assessed by visual inspection of the chromatograms. Alignments were generated with Clustal X v. 2 [38] under the default settings and were trimmed to remove part of the 3’ end, for which many sequences were incomplete. Point substitutions that caused stop codons or nonconservative changes in amino acid were checked against the original sequencing trace files. In some cases, the sequence was eliminated from further analysis at this stage. When necessary, the alignments obtained were then improved manually using the program MEGA v. 7.0 [37].

The phylogenetic reconstruction was based on parsimony (MP) [39], maximum likelihood (ML) [40–41], and Bayesian inference (BI) [42] methods. In all analyses, gaps were treated as missing data.

Parsimony analyses were performed using TNT ver. 1.1 [43] with Fitch parsimony [39] as the optimality criterion. All characters were equally weighted and treated as unordered. A heuristic search was conducted using 1000 random taxon-addition replicates, with the tree-bisection-reconnection (TBR) algorithm, saving up to 15 trees per replicate to prevent extensive swapping on islands with many trees. The resulting trees were then used as starting trees for a second-round search using TBR branch swapping with an upper limit of 10,000 trees. Nonparametric bootstrap support (BS) was estimated using 10,000 pseudo-replicates, and the same parameters were used in our MP analyses [44]. Bootstrap percentages of 50 to 80 were considered weak, 81 to 90 moderate, and > 90 strong.

ML analyses were conducted using RAxML-HPC2 on XSEDE (v. 8.2.12) [45] in the Cyberinfrastructure for Phylogenetic Research (CIPRES) Portal v. 3.3 [46]. For this analysis we used the implemented algorithm, which allows one to perform optimal tree searches and obtain bootstrap support [44] in one single analysis [47]. To this end, we performed 1000 bootstrap replicates with a subsequent search of the maximum likelihood tree, using the GTRGAMMA nucleotide substitution model [45], individual per-site substitution rates (-c), and default setting of likelihood acceptance (-e), 25 and 0.1, respectively. Bootstrap percentages of 50 to 80 were considered weak, 81 to 90 moderate, and > 90 strong.

Bayesian analyses were performing using MrBayes v. 3.2.7a [48] in the CIPRES Portal [46]. To determinate the best-fitting nucleotide substitution model, data were submitted to jModeltest 2.1.1 [49] and the Akaike information criterion (AIC) selected TVM+I+G. The dataset was analyzed in two independent runs of 10 million generations, each with four Markov chains (one cold and three heated chains), sampling every 1000 generations. Convergence of the runs was assessed by checking the status of parameters in Tracer v.1.7 [50] to ensure the stationarity of each run. Likelihoods of the trees produced by each run were analyzed graphically using Tracer v.1.7 [50] and, after discarding the initial 2500 trees of each run as burn-in (25%), the remaining trees (15,002) were used to generate a 50% majority-rule consensus tree. The cutoff for strong support in the Bayesian analyses was 0.95 (roughly equal to p < 0.05) posterior probabilities and values below 0.8 were considered not supported.

## Results

The aligned data matrix for 178 accessions consists of 2077 nucleotide positions, of which 273 characters were phylogenetically informative. The parsimony analyses found 40 trees of 768 steps (uninformative characters excluded), with a consistency index (CI) of 0.464 and a retention index (RI) of 0.802. The strict consensus tree from MP, the Bayesian 50% majority-rule consensus tree, and the ML tree all produced similar topologies showing the same strongly supported clades; thus, only the BI tree is presented here, along with branch support obtained under MP and ML analyses (Fig 1). The aligned data matrix and trees from the three methods of analysis are available at Repositorio Institucional CONICET Digital [51]: http://hdl.handle.net/11336/163438.

**Fig 1.**
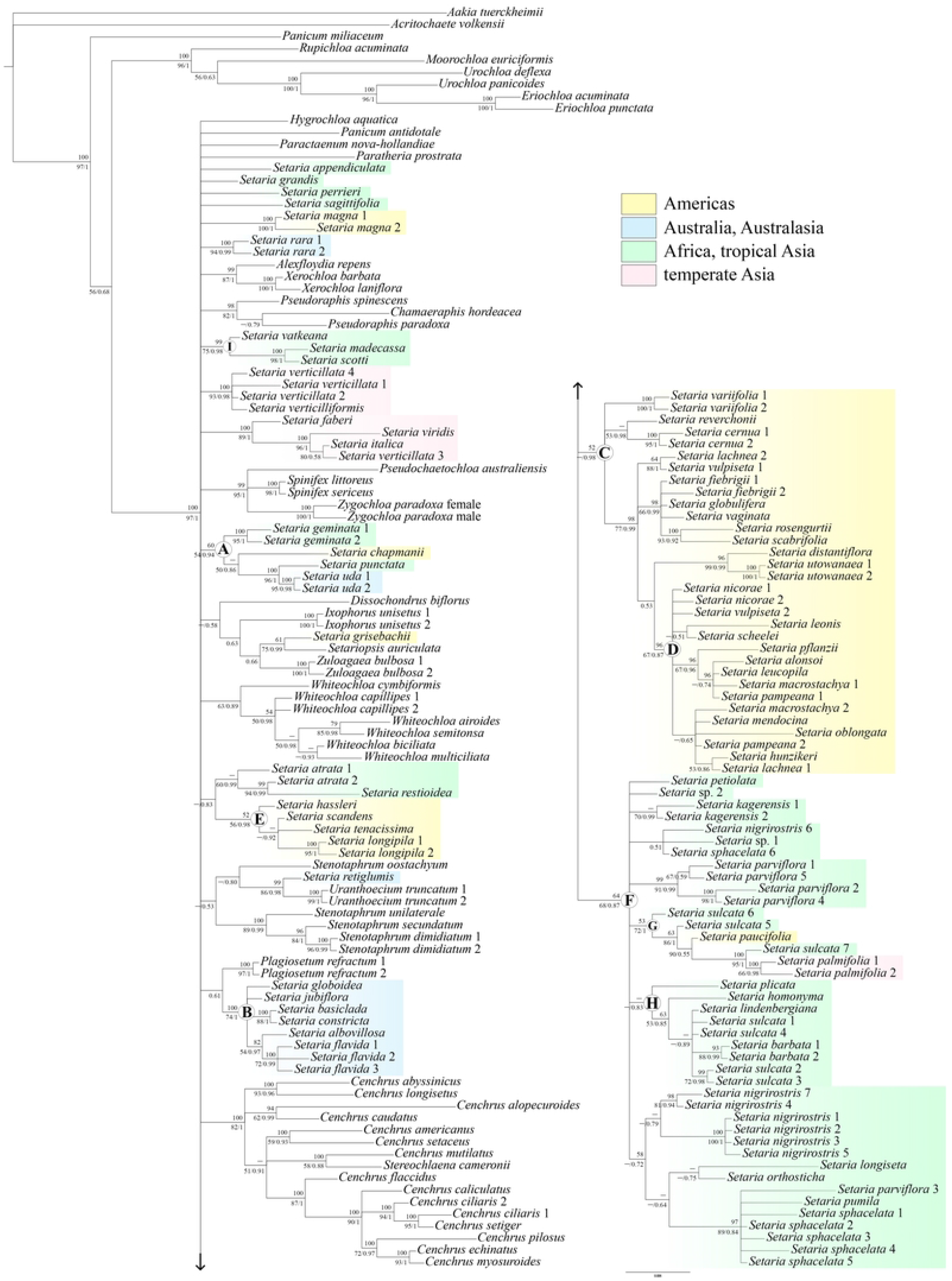
Phylogenetic reconstruction of *Setaria* within the subtribe Cenchrinae, based on the chloroplast *ndh*F gene. Bootstrap supports from parsimony are listed above the branches, and bootstrap supports from maximum likelihood / posterior probabilities from Bayesian inference are listed below the branches. Nodes with “–” have bootstrap supports < 50% and the clades denoted by letters are discussed in the text. Accepted names for *Setaria* species follow [2, 5, 22] and the remainder Cenchrinae taxa plus the outgroup follow [23].

For ten taxa we were able to add a second voucher (i.e., *Plagiosetum refractum* (F. Muell.) Benth., *Setaria atrata* Hack., *S. cernua, Setaria geminata* (Forssk.) Veldkamp, *Setaria magna* Griseb., *Setaria nicorae* Pensiero, *Setaria palmifolia* (J. Koenig) Stapf, *Setaria rara* (R. Br.) R.D. Webster, *Setaria uda* (S.T. Blake) R.D. Webster, and *Uranthoecium truncatum* (Maiden & Betche) Stapf), as well as include two accessions for *Setaria longipila* E. Fourn., *Setaria utowanaea* (Scribn.) Pilg., *Setaria variifolia* (Swallen) Davidse, *Stenotaphrum dimidiatum* (L.) Brongn., and *Whiteochloa capillipes* (Benth.) Lazarides, four new accessions for the polymorphic *Setaria sulcata* Raddi (vouchers 4–7), and one more accession for *Setaria nigrirostris* (Nees) T. Durand & Schinz (voucher 7) (Table 1; Fig 1). With the exception of *S. nicorae* (subclade D) and *W. capillipes* (“Whiteochloa clade”) whose positions are uncertain, in most cases the two accessions of the same species had identical or nearly identical sequences and were placed together by the three analyses. The two accessions of *S. atrata* were separated in the tree but still formed a clade, and in *S. sulcata* mutations in the sequences led the accessions to distinct placements (i.e., separated into two clades) (Fig 1).

Based on the phylogenetic evidence presented here, the subtribe Cenchrinae sensu [11–12] is paraphyletic, in agreement with previous results [10, 30, 52]. Our analyses also showed *Setaria* to be polyphyletic, with its species distributed in at least ten distinct clades (Fig 1). The relationships among clades remains largely unresolved but all combined analyses placed the 32 new accessions of *Setaria* in six clades and three subclades (A–I; Fig 1), as detailed next.

Clades A and B group most of the *Setaria* species considered in the past as *Paspalidium* [i.e., *Setaria albovillosa* (S.T. Blake) R.D. Webster, *Setaria basiclada* (Hughes) R.D. Webster, *Setaria chapmanii* (Vasey) Pilg., *Setaria constricta* (Domin) R.D. Webster, *Setaria flavida* (Retz.) Veldkamp, *Setaria geminata, Setaria. globoidea* (Domin) R.D. Webster, *Setaria jubiflora* (Trin.) R.D. Webster, *Setaria punctata* (Burm. f.) Veldkamp, and *S. uda*]. Within clade A (Bayesian posterior probability (BPP) 0.94 / ML bootstrap (MLB) 54 / parsimony bootstrap (PB) 60), the two accessions of *S. uda* (BPP 0.98 / MLB 95 / PB 100) are sister to *S. punctata* (BPP 1 / MLB 96 / PB 100), and both are related to the American *S. chapmanii* (BPP 0.86 / MLB 50 / PB < 50). The two accessions of *S. geminata* (BPP 1 / MLB 95 / PB 100) are strongly (BI) supported as sister to these species.

Clade B, strongly (BI and MP) supported (BPP 1 / MLB 74 / PB 100), consists of species of *Setaria* native to Australia and Australasian regions. Although the relationships within this clade are not clearly defined, *S. basiclada* was sister to *S. constricta* (BPP 1 / MLB 88 / PB 100), as well as *S. albovillosa* resolved as sister to the three accessions of *S. flavida* (BPP 0.97 PP / MLB 54 / PB 82). The two accessions of *Plagiosetum refractum* (BPP 1 / MLB 97 / PB 100) were resolved as sister group to all clade B species with a weak branch support in the BI analysis (BPP 0.61).

Clades C–E include most of the American *Setaria* species. Within clade C (BPP 0.98 / MLB < 50 / PB 52), the two accessions of *S. variifolia* were consistently placed together (BPP 1 / MLB 100 / PB 100), as well as the two accessions of *S. cernua* (BPP 1 / MLB 95 / PB 100) closely related to *Setaria reverchonii* (Vasey) Pilg. (BPP 0.98 / MLB 53 / PB < 50), although their positions within this clade are unclear. *Setaria leonis* (Ekman ex Hitchc.) León were resolved closely related to *Setaria scheelei* (Steud.) Hitchc. with almost no support in BI and ML analyses (BPP 0.51 / MLB < 50) and not recovered in MP. The two accessions of *S. nicorae* remain unresolved, forming part of a polytomy within subclade D strongly supported by MP analysis (BPP 0.87 / MLB < 50 / PB 96). Bayesian and ML approaches also indicated a close relationship between *Setaria hunzikeri* Anton and *Setaria lachnea* (Nees) Kunth (1), with a moderate (BI) branch support (BPP 0.86 / MLB 53), but this relationship was not recovered in MP. The two accessions of *S. utowanaea* (BPP 1 / MLB 100 / PB 100) are sister to *Setaria distantiflora* (A. Rich.) Pilg. (BPP 0.99 / MLB 99 / PB 96), and both were resolved as sister clade to all species of subclade D.

Clade E was strongly supported only by the BI analysis (BPP 0.98 / MLB 56 / PB 52) and its members are not related to the species of the larger American clade. It includes *Setaria hassleri* Hack. sister to *Setaria scandens* Schrad., *Setaria tenacissima* Schrad., and the two accessions of *S. longipila* (BPP 1 / MLB 96 / PB 100), which are grouped in a polytomy (BPP 0.92 / MLB < 50 / PB < 50). *Setaria restioidea* (Franch.) Stapf is nested with one accession of *S. atrata* (BPP 0.99 / MLB 94 / PB 99), and both are related to *S. atrata* 2 (BPP 0.99 / MLB 60 / PB < 50). This clade was resolved as sister to all species of clade E (BPP 0.83 / MLB < 50), relationship that was not supported in MP.

Clade F is moderately (BI) supported (BPP 0.87 / MLB 68 / PB 64) and groups most of the African *Setaria* species. The two accessions of *S. palmifolia* were placed together (BPP 0.98 / MLB 66 / PB 100) in subclade G with *S. sulcata* 7, *Setaria paucifolia* (Morong) Lindm., and *S. sulcata* 5 and 6 supported as its successive sisters (BPP 1 / MLB 95 / PB 100; BPP 0.55 / MLB 90; BPP 1 / MLB 86 / PB 63; and BPP 1 / MLB 72 / PB 53, respectively). The other four accessions of *S. sulcata* (1–4) were resolved in a polytomy (BPP 0.89 / MLB < 50 / PB < 50) within subclade H together with the two accessions of *Setaria barbata* (Lam.) Kunth (BPP 0.99 / MLB 88 / PB 93) and *Setaria lindenbergiana* (Nees) Stapf., *Setaria homonyma* (Steud.) Chiov. and *Setaria plicata* (Lam.) T. Cooke were resolved as successive sisters to all subclade H species with a moderate branch support in the BI analysis (BPP 0.85 / MLB 53 / PB 63; BPP 0.83 / MLB < 50 / PB < 50, respectively).

Clade I groups three *Setaria* species restricted to the Madagascar archipelago in a strongly supported clade by BI and ML analyses (BPP 0.98 / MLB 87 / PB 99). Within this clade, *Setaria madecassa* A. Camus is sister to *Setaria scottii* (Hack.) A. Camus (BPP 1 / MLB 98 / PB 100), and both are related to *Setaria vatkeana* K. Schum.

*Setaria perrieri* A. Camus, *Setaria sagittifolia* (A. Rich.) Walp., *S. magna*, and *S. rara* are the only four species that have not been consistently assigned/related to any of the retrieved clades and, consequently, their positions are uncertain. Parsimony and ML analyzes found weak support (bootstrap < 50) for a sister relationship between *S. perrieri* with “Cenchrus clade” but this grouping was not supported in BI. Although a second voucher of *S. magna* and *S. rara* were included here, and the two accessions of each species were strongly supported as sisters (BPP 1 / MLB 100 / PB 100; BPP 0.99 / MLB 94 / PB 100, respectively), their placements in this phylogeny remain unclear.

## Discussion

### Relationships within *Setaria* and major results

Here the analyzed species of *Setaria* were recovered as a set of at least ten unrelated groups, consisting mostly of several phylogenetically disparate clades distributed in warm regions around the world (Fig 1). In addition, as *Setaria* lacks unique distinctive characters, it will need to be expanded to include some new elements when a solid phylogeny becomes available. Relationships among species were similar to those shown in previous *ndh*F-based phylogenies [13–14] with notable differences mainly in the composition of the previously proposed American clades. Here, most of the American *Setaria* species were resolved in two main clades, one major (clade C, corresponding to clade X of [14]) and one minor (clade E, corresponding to clade II of [14]), both morphologically quite distinct. The major clade was originally composed of South American perennial species; however, due to the placement of some species of the subgenera *Paurochaetium* and *Reverchoniae* (Table 2) within this clade, its range was extended to Central and North America. Our phylogeny also identified a clade related to the morphology of the species rather than its geographic origins (clade A), which groups *S. chapmanii*, a taxon previously treated in subgenus *Paurochaetium* [3] (Fig 1).

As expected, the subgenera *Paurochaetium* and *Reverchoniae* (Table 2) are non-monophyletic (Fig 1) like the other subgenera of *Setaria*. Although they share morphological similarities, five of the six species analyzed (i.e., *S. distantiflora, S. leonis, S. reverchonii, S. variifolia*, and *S. utowanaea*) were resolved within the major American clade, according to their geographic origins. Except for *S. distantiflora* sister to *S. utowanaea*, our analyses did not place *S. leonis, S. reverchonii*, and *S. variifolia* together, indicating a more distant relationship among them. On the other hand, *S. chapmanii*, also an American species, was unambiguously strongly supported within clade A, related to species with inflorescences “Paspalidium type”. While the previously recognized subgenera *Paurochaetium* and *Reverchoniae* fail to define monophyletic groups in *Setaria*, they are useful as a way to organize the discussion about relationships of the studied species. An analysis of the different clades and relationships among species are discussed next.

### Relationships of taxa added in this study

As mentioned earlier, the taxonomic history of subgenera *Paurochaetium* and *Reverchoniae* are linked that of the genus *Paspalidium*. Species of subgenus *Paurochaetium* were originally described as a subgenus of *Panicum* [33] to accommodate taxa in which setae are present only at the ends of the primary branches of the inflorescence. Subgenus *Paurochaetium* was first placed under *Setaria* at the rank of section by [53], and elevated to subgenus by [3]. Following [33]’s concept, [17] established the genus *Paspalidium*, segregating it from *Setari*a but, species of subgenus *Paurochaetium* were transferred to *Paspalidium* only decades later [34]. As the circumscriptions of the two genera overlap and the distinction between them is somewhat arbitrary, the *Paspalidium* species were transferred back to *Setaria* ([1, 19–20]; see Table 2 for a synopsis of species and different classifications for the taxa of subgenera *Paurochaetium* and *Reverchoniae*), a result supported by molecular analyses [10, 13–14].

Subgenera *Paurochaetium* (five species) and *Reverchoniae* (two species) (Table 2; [5, 18]) include caespitose perennial plants distributed from the United States (New Mexico, West South Central, and Florida) to the north of South America (Venezuela and Colombia), being mostly concentrated in the Caribbean [3, 5, 18, 23, 34]. *Setaria chapmanii*, analyzed here for the first time, grows on limestone, coral, shell or sandy soils in Florida Keys, the Bahamas, Cuba, and the Yucatan Peninsula [3]; its panicles have branches with spikelets biseriate, the blunt first glume turned away from the rachis and the back of the upper lemma toward it, and a single bristle present below the terminal spikelets [34]. Although this species was previously treated within the subg. *Paurochaetium* [3], the well-ordered arrangement of its spikelets in unilateral spikes is highly anomalous in this group [34], as well as the lack of the lower palea [3]. In our phylogeny, *S. chapmanii* is placed in clade A and turned out to be the only species of subg. *Paurochaetium* that is related to the others previously considered in *Paspalidium* (the remaining species of subg. *Paurochaetium* were included in clade C).

Clade A groups species with inflorescences “Paspalidium type” related to wet/aquatic habitats and, except for *S. chapmanii* which have slender culms, its members are characterized by having spongy culms [2, 34, 54]. The relationship between *S. chapmanii* and *S. geminata*, for sharing the same type of environment, has been previously highlighted [34]. *Setaria geminata* is native to Africa and Asia, introduced unintentionally in tropical and subtropical areas of other continents [2]. It is an aquatic species with thick and spongy culms, while *S. chapmanii* inhabits temporary pools and marshes, and is characterized by having culms mostly simple, erect, slender, and smooth [34, 54]. The spongy culms of *S. geminata* are also shared with *S. punctata* and *S. uda*. From the former species *S. geminata* is distinguished by having spikelets ovoid and lower palea well-developed while in *S. punctata* the spikelets are ellipsoid and lack lower palea [2]. *Setaria uda* is a species native to Australia and Papua New Guinea [2] and its position within clade A is confirmed here by the addition of a second voucher (i.e., *S. uda* 2). It differs from *S. geminata* and *S. punctata* mainly by having caespitose habit; it lacks rhizomes and has smaller spikelets [2].

In [14], *S. magna, S. rara* and *Plagiosetum refractum* were resolved as successive sisters to the “Paspalidium clade” (i.e., clade B), relationships not retrieved by our analyses (with exception of *Plagiosetum refractum* whose position is poorly supported only by BI). *Setaria rara* is endemic to Australia, commonly found in arid areas associated with creeks or lagoons [2]. It was previously included in *Paspalidium* and ML analysis suggested a sister relationship between *S. rara* and clade B species, although without support (bootstrap < 50). *Setaria rara* is morphologically similar to *S. basiclada*, in that is shares an annual habit [2]; however, its position remains unresolved even with the addition of a second voucher. *Setaria magna* is also an annual species but it is native to tropical and subtropical Americas and is morphologically different from *Paspalidium*. It is distinguished from other species of *Setaria* by its robust aspect with culms as much as 4 m tall and ligules forming an inverted “V” [5]. Its placement is not yet defined; however, our analyses corroborated that *S. magna* is not related to the American clades, and suggests a more distant relationship with clade B species.

Clade C groups most of the American perennial species of *Setaria* and, as in previous studies [13–14], it was retrieved in all analyses. *Setaria cernua*, whose position was unclear in [14], was consistently supported within this clade, nested with *S. reverchonii*. It is characterized by having conspicuous superficial rhizomes, tillers with strongly keeled leaves resembling those of some Iridaceae, lower anthecium male with developed anthers, and upper anthecium shorter than the spikelet [5]. This unique combination of characters states led [5] to establish the monotypic subg. *Cernatum*, which was not supported by our findings, and also disagrees with previous results [14] which had recovered it in an isolated position.

The two species of subg. *Reverchoniae* (Table 2) were also placed in clade C but they do not appear to be related to each other nor to the subg. *Paurochaetium* taxa. Subgenus *Reverchoniae* was erected to accommodate species with panicles erect, spikelets randomly disposed on the branches, and the central inflorescence axis scabrous [18]. *Setaria variifolia* differs from *S. reverchonii* mainly in having the lower palea well-developed and by the geographic distribution in the Yucatán peninsula of Mexico and Mesoamerica (vs. Texas, New Mexico, and Oklahoma (United States) and northern Mexico) [18]; its placement within the larger American clade is confirmed here by sequencing of two vouchers, but its relationships remain unknown.

Within subclade D, *S. leonis* was resolved in a weakly supported position sister to *S. scheelei*, a unexpected result as these species are morphologically very different from each other and do not grow sympatrically. *Setaria leonis* is endemic to the Caribbean islands, commonly found on rocky slopes and clearings while *S. scheelei* is native to southwest and south-central United States to Mexico and prefers shaded habitats on alluvial soils of limestone canyons and river bottoms [3, 23]. *Setaria leonis* shares the slender culms, geographic distribution and habitats with *Setaria pradana* (León ex Hitchc.) León [3]; however, we were not able to analyze the latter species because the *ndh*F failed in all amplification attempts.

*Setaria scheelei* has been assigned to subg. *Setaria* [3] and was included in this analysis since it shares a geographic distribution pattern similar to that of subg. *Paurochaetium*. It is a highly polymorphic species characterized by having robust aspect, culms usually geniculate at the base, blades usually flat and pubescent, and the upper lemma short-apiculate, incurved, finely cross-wrinkled [3]. It is morphologically similar to *Setaria macrostachya* Kunth but our data suggest a more distant relationship between them, although the position of the latter species in the tree is unclear.

*Setaria hunzikeri*, here analyzed for the first time, was resolved sister to one accession of *S. lachnea*, also within subclade D. The two species are important forage grasses native in South America and are morphologically very similar [5], so their close relationship in our analyses is not surprising. *Setaria hunzikeri* differs from *S. lachnea* by having hirsute and narrower blades and smaller inflorescences up to 8 cm long (vs. blades glabrous or scabrous and inflorescences ranging from 7 to 25 cm long) [5].

*Setaria nicorae* was represented in [14]’s phylogeny by a partial sequence and it was placed in a polytomy together with other South American perennial species. Here, by including a second voucher with a complete *ndh*F sequence, we confirmed the placement of *S*.*a nicorae* within the major American clade but its relationships remain unknown. Morphological similarities between *S. nicorae* and *S. utowanaea* have been noted by [5], mainly by sharing the caespitose habit with conspicuous rhizomes, spikelets ovoid, and the upper glume 5–7-nerved; however, our analyses indicated a more distant relationship between them. The latter species is sister to *S. distantiflora* and both were resolved as sister to all subclade D species but this relationship was recovered only by the BI analysis and with almost no support. *Setaria distantiflora* and *S. utowanaea* are commonly found in open, rocky soils; they are morphologically distinct but similar in general aspect. *Setaria distantiflora* is endemic to the Caribbean and is characterized by a caespitose habit, lacking conspicuous rhizomes, ligules as a fringe of very short hairs, and spikelets lanceolate-ellipsoid while *S. utowanaea* has short rhizomes, ligules membranous-ciliate, spikelets ovoid and is more widely distributed (i.e., Caribbean, Colombia to Venezuela) [3, 5].

As presented by [14], the minor American clade (clade E) groups annual species with “bottle-brush inflorescences” (i.e., cylindrical, dense, and continuous spiciform panicles), and both antrorse and retrorse prickles on the same bristle, the latter indicated as potential morphological synapomorphy of this clade [14]. In this phylogeny, bootstrap supports for relationships of clade E are weaker than that retrieved previously, possibly because of the placement of *S. longipila*, here analyzed for the first time, within this clade. *Setaria longipila* is also an annual species but, its subspiciform panicles [3] are distinctive. On the other hand, *Setaria grisebachii* E. Fourn., another annual species with inflorescences similar to those of *S. longipila*, is sister to *Setariopsis auriculata* (E. Fourn.) Scribn. (BPP 0.99 / MLB 75 / PB 61), and are both related to *Zuloagaea bulbosa* (Kunth) E. Bess. The position of *S. grisebachii* outside clade E will have to be verified by inclusion of multiple accessions and other morphologically similar American species (e.g., *Setaria liebmanni* E. Fourn.) in further analyses.

Most African *Setaria* species are grouped in clade F but no obvious morphological characters shared by all members were identified. In [14]’ phylogeny, *S. atrata* was represented by a partial sequence and weakly supported as sister to *S. restioidea* and *S. paucifolia* (Morong) Lindm. Here, by including a second voucher of *S. atrata* with a complete sequence, we have confirmed the sister relationship with *S. restioidea*, although its two accessions were not resolved together; however, our data indicated a more distant relationship among these two species with *S. paucifolia*, as the latter was resolved within subclade G. *Setaria atrata* and *S. restioidea* are found in swampy places, on clay and saline soils, and their close relationship and morphological similarities have been previously discussed [2, 14]. *Setaria atrata* is distinguished by having the upper anthecium strongly papillose with transverse wrinkles and lower lemma membranous while *S. restioidea* has the upper anthecium smooth, shiny, finely papillose, lacking prominent wrinkles, and lower lemma coriaceous [2]. The convolute and rigid blades are also shared with the South American *S. paucifolia* but our findings suggest that the latter species is related to *S. sulcata* and *S. palmifolia*, which have plicate blades. Although this result is unexpected, it seems that species of subclade G share not morphology but rather habitats (i.e., they are frequent in moist and shady places, streambanks, and along forest paths [2, 5]). As we were not able to analyze multiple vouchers of *S. paucifolia*, the question on the possible Africa–South America disjunction noted by [14] remains unanswered.

The second group with blades plicate is represented in subclade H and consists of an intricate polymorphic species complex with chromosome counts ranging from 2*n*=32 to 2*n*=56, and *n*=16 and18 [55–63]. Several taxa with plicate blades have been synonymized under the name *S. sulcata* due to its apparent substantial plasticity and overlapping of morphological limits and in geographic distribution [2, 5]. Our sampling included specimens determined as *S. sulcata* (voucher 1), *Setaria poiretiana* (Schult.) Kunth (vouchers 2–5), and *Setaria megaphylla* (Steud.) T. Durand & Schinz (vouchers 6 and 7). Although our analysis is not conclusive regarding the taxonomic position of this species complex, the distinct placements of *S. sulcata* in the tree did not support the recognition of them as a single widespread species, in disagreement with previous results [14].

Clade I represents a segregate lineage grouping three species endemic to Madagascar, as suggested in earlier studies [30]. *Setaria madecassa* and *S. scottii* are found on granite or basaltic soils of sub-humid savannas [2]. The former species is characterized by having annual caespitose habit with culms geniculate, panicle open with ascending branches and a single seta below each spikelet, while *S. scottii* includes perennial plants, with culms decumbent, panicles contracted, and gibbous spikelets accompanied by a short seta or lacking a seta [2]. *Setaria vatkeana* is an annual species of forested humid areas, differing by its culms erect and unbranched, blades pseudopetiolate, sulcate spikelets with a single seta on all of spikelets, and lower lemma indurated, coriaceus and papillose [2]. *Setaria perrieri* is another endemic species to Madagascar but its position in this phylogeny remains unresolved. Its weakly supported sister relation to the “Cenchrus clade” in MP and ML should be considered provisional until confirmed by additional genes and accessions.

### “Cenchrus clade”

*Cenchrus* L. is a cosmopolitan genus with approximately 120 species [52] characterized by having one or several spikelets accompanied by one bristle or surrounded by an involucre of multiple bristles, or with bristles fused in a cup-like structure [64]. It is monophyletic only when *Pennisetum* Rich. and the monotypic *Odontelytrum* Hack. are included within it; however, recent molecular phylogenetic studies [30] and our findings showed *Cenchrus* paraphyletic with *Stereochlaena cameronii* (Stapf) Pilg. embedded in it. *Stereochlaena cameronii* is morphologically quite distinct from *Cenchrus* in having digitate racemes, imbricate paired spikelets, and lower lemma awned [23]. Therefore, to reach any decision on the inclusion of this species within *Cenchrus* its placement in the tree must be confirmed by additional accessions and more variable markers.

*Pseudochaetochloa australiensis* Hitchc., an endemic species to Australia, is considered as a synonym of *Cenchrus arnhemicus* (F. Muell.) Morrone in [23]; however, this treatment is not supported by our analyses. Here, *Pseudochaetochloa australiensis* forms a strongly supported clade with the dioecious Australian *Spinifex* and *Zygochloa*, which corroborates its classification as an independent genus of *Cenchrus. Pseudochaetochloa australiensis* is distinguished from these two by having monoecious 2-flowered spikelets bearing a single bristle subtending many of the spikelets, lower anthecium well developed, and both lemmas membranous, similar in size, shape, and texture [65].

### *Stenotaphrum* Trin

*Stenotaphrum* is a primarily tropical genus including seven species [23, 66–67] and, as in *Paspalidium*, its secondary-order inflorescence branches end in a bristle [67]. The placement of *Stenotaphrum* within subtribe Cenchrinae and its phylogenetic relationships have been uncertain due to limited data from previous studies (i.e., in [14] it was represented only by *Stenotaphrum secundatum* (Walter) Kuntze). Here, by increasing the number of species sampled (total of four), our results corroborate the close relationship of *Stenotaphrum* with *Setaria retiglumis* (Domin) R.D. Webster (syn. *Paspalidium retiglume* (Domin) Hughes) and *Uranthoecium truncatum*, retrieving it as paraphyletic, as suggested by [30]. *Stenotaphrum* is distinguished from *Setaria, Paspalidium* and the monotypic *Uranthoecium* by having the main inflorescences axis thickened and flattened, with the secondary branches embedded in it [66–67]. *Uranthoecium truncatum* is characterized by having short lateral branches, disarticulating rachis and truncate glumes, a set of features unique in this clade. As in previous analyses [14], *Uranthoecium truncatum* was strongly (BI and MP) supported as related to *S. retiglumis*. Both are caespitose annual species endemic to Australia and exhibit a very similar foliar anatomy [2, 21], although *S. retiglumis* is morphologically more similar to the other *Setaria* species.

### *Acritochaete volkensii* Pilg

The monotypic *Acritochaete volkensii* is an annual species found in shady forests of tropical Africa [23, 68], currently treated within the subtribe Cenchrinae [11–12]. It has inflorescences bearing persistent setae, a character state shared with the “bristle clade”; however, our results placed *Acritochaete volkensii* outside of the Cenchrinae, as also indicated in [30, 52]. Despite the morphological similarities with Cenchrinae, the photosynthetic C_3_ pathway of *Acritochaete volkensii* is unusual within this subtribe, which includes C_4_ NADP–ME plants [10, 12]. According to [30], *Acritochaete* Pilg. is closely related to members of the Boivinellinae Pilg., subtribe known as “the forest shade clade” [25, 30] and which groups mostly physiologically C_3_ genera [10, 12]. Thus, for both subtribes to be monophyletic *Acritochaete volkensii* must be recognized within the subtribe Boivinellinae.

### Final considerations on the general approach and needs for future studies in *Setaria*

This article added the missing information on the knowledge of the relationships of the subgenera *Paurochaetium* and *Reverchoniae*. Although both were not recovered as natural groups, as the other infrageneric categories of *Setaria*, this step was required since its species had never been analyzed in any previous molecular phylogeny. Here, we were able to clarify many of the pending questions and identify clades morphologically distinctive not necessarily correlated with the geographic origin of species as proposed earlier. On the other hand, this paper also shows the limits of what can be done with a single gene in *Setaria* and Cenchrinae: there are 178 taxa and 273 informative characters, which were not enough to resolve the tree strongly supported. For estimating bootstrap support, as a general guide, every branch requires at least three mutations to have confidence in phylogenetic analyses [44]. The problem of too few variable sites affects the MP and ML analyses in particular and may partly explain the low bootstrap values, since there are not enough characters. Because Bayesian approaches estimate support in a different way, they are less susceptible to the limitation of too few characters. Here, the branches with BPP values of 0.98 but low MLB or PB were supported by only one or two characters (e.g., clades C and E). As we have completed the *ndh*F-based phylogeny for all existing infrageneric categories within *Setaria*, the next step is for future studies to use markers that allow capturing more mutations, such as other chloroplast protein-coding genes or whole plastomes [69–71], and/or low-copy nuclear genes (LCNGs) [72–73]. As it is well-known that chloroplast genome cannot recover reticulations caused by allopolyploids and that plastomes phylogenies give an incomplete picture of the history of any group with hybridization [74–75], the results should be confirmed by studies that include LCNGs, which hold great potential to improve the robustness of phylogenetic trees [76], and therefore may be a key to generating a better understanding of the complex relationships in *Setaria*.

## Acknowledgments

The authors thank to Dr. Fernando O. Zuloaga (Instituto de Botánica Darwinion), for his assistance during the stages of this study, and to Dr. Elizabeth A. Kellogg (Donald Danforth Plant Science Center), who kindly read our manuscript, for her comments and suggestions that greatly strengthened the final version.

## References

1. Webster RD. Nomenclature of Setaria (Poaceae: Paniceae). Sida 1993; 15(3): 447–489.

2. Morrone O, Aliscioni SS, Veldkamp JF, Pensiero JF, Zuloaga FO, Kellogg EA. Revision of the Old World Species of Setaria (Poaceae: Panicoideae: Paniceae). Syst. Bot. Monogr. 2014; 96: 01–161.

3. Rominger JM. Taxonomy of Setaria (Gramineae) in North America. Illinois Biol. Monogr. 1962; 29: 01–132.

4. Prasada Rao KE, Wet JMJ, Brink DE, Mengesha MH. Intraspecific variation and systematics of cultivated Setaria italica, foxtail millet (Poaceae). Econ. Bot. 1987; 41: 108–116.

5. Pensiero JF. Las especies sudamericanas del género Setaria (Poaceae, Paniceae). Darwiniana. 1999; 37(1–2): 37–151.

6. Stapf O, Hubbard OE. Setaria P. Beauv. In: Oliver D, editor, Flora of tropical Africa, Gramineae (Maydeae-Paniceae), vol. 9. Ashford: Reeve & Co; 1930. pp. 768–866.

7. Callmander MW. Catalogue of the Vascular Plants of Madagascar (Madagascar Catalogue), Missouri Botanical Garden and Antananarivo [online]. Available from: http://legacy.tropicos.org/Project/Madagascar [accessed 10 February 2022].

8. BFG (The Brazil Flora Group). Brazilian Flora 2020: Leveraging the power of a collaborative scientific network. Taxon. 2021; 00: 01–21. DOI: https://doi.org/10.1002/tax.12640.

9. Sousa VF, Santos CAG, Boldrini II. Setaria in Flora e Funga do Brasil, Jardim Botânico do Rio de Janeiro [online]. Available from: http://floradobrasil.jbrj.gov.br/reflora/floradobrasil/FB13581 [accessed 10 February 2022].

10. Morrone O, Aagesen L, Scataglini MA, Salariato DL, Denham SS, Chemisquy MA, et al. Phylogeny of the Paniceae (Poaceae: Panicoideae): integrating plastid DNA sequences and morphology into a new classification. Cladistics. 2012; 28(4): 01–24.

11. Soreng RJ, Peterson PM, Romaschenko K, Davidse G, Zuloaga FO, Judziewicz EJ, et al. A worldwide phylogenetic classification of the Poaceae (Gramineae). J. Syst. Evol. 2015; 53(2): 117–137.

12. Soreng RJ, Peterson PM, Zuloaga FO, Romaschenko K, Clark LG, Teisher JK, et al. A worldwide phylogenetic classification of the Poaceae (Gramineae) III: An update. J. Syst. Evol. 2022; 60(3): 476–521.

13. Kellogg EA. Evolution of Setaria. In: Doust A, Diao X, editors, Genetics and Genomics of Setaria, Plant Genetics and Genomics: Crops and Models, vol. 19. NewYork: Springer, Cham, Swiss; 2017. pp. 01–27.

14. Kellogg EA, Aliscioni SS, Morrone O, Pensiero J, Zuloaga FO. A phylogeny of Setaria (Poaceae, Panicoideae, Paniceae) and related genera based on the chloroplast gene ndhF. Int. J. Plant Sci. 2009; 170(1): 117–131.

15. GPWG II (Grass Phylogeny Working Group II). New grass phylogeny resolves deep evolutionary relationships and discovers C_4_ origins. New. Phytol. 2012; 193(2): 304–312.

16. Clayton WD, Renvoize SA. Genera Graminum. Grasses of the World. Kew. Bull., Addit. Ser. 1986; 13: 01–389.

17. Stapf O. Paspalidium. In: Prain D, editor, Flora of tropical Africa, Gramineae (Maydeae-Paniceae), Ashford: Reeve & Co.; 1920. pp. 582–586.

18. Fox WE, Hatch SL. New combinations in Setaria (Poaceae: Paniceae). Sida. 1999; 18(4): 1037–1047.

19. Veldkamp JF. Miscellaneous notes on Southeast Asian Gramineae. IX. Setaria and Paspalidium. Blumea. 1994; 39(1/2): 373–384.

20. Webster RD. Nomenclatural changes in Setaria and Paspalidium (Poaceae: Paniceae). Sida. 1995; 16(3): 439–446.

21. Aliscioni SS, Ospina JC, Gomiz NE. Morphology and leaf anatomy of Setaria s.l. (Poaceae: Panicoideae: Paniceae) and its taxonomic significance. Plant. Syst. Evol. 2016; 302: 173–185.

22. Pensiero JF. Setaria P. Beauv. In: Zuloaga FO, Morrone O, Davidse G, Filgueiras TS, Peterson PM, et al., editors, Catalogue of New World Grasses (Poaceae): III. Subfamilies Panicoideae, Aristidoideae, Arundinoideae, and Danthonioideae. Washington, D.C.: Smithsonian Institution; 2003. pp. 46: 569–593.

23. POWO (Plants of the World Online). Facilitated by the Royal Botanic Gardens, Kew [online]. Available from: http://www.plantsoftheworldonline.org/ [accessed 8 March 2022].

24. Thiers B. Index Herbariorum: A Global Directory of Public Herbaria and Associated Staff, New York Botanical Garden’s Virtual Herbarium, continuously updated [online]. Available from: http://sweetgum.nybg.org/science/ih/ [accessed 8 March 2022].

25. Giussani LM, Cota-Sánchez JH, Zuloaga FO, Kellogg EA. A molecular phylogeny of the grass subfamily Panicoideae (Poaceae) shows multiple origins of C_4_ photosynthesis. Amer. J. Bot. 2001; 88(11): 1993–2012.

26. Doust AN, Kellogg EA. Inflorescence diversification in the panicoid ‘‘bristle grass’’ clade (Paniceae, Poaceae): evidence from molecular phylogenies and developmental morphology. Am. J. Bot. 2002; 89(8): 1203–1222.

27. Doust AN, Penly AM, Jacobs SWL, Kellogg A. Congruence, conflict and polyploidization shown by nuclear and chloroplast markers in the monophyletic bristle clade (Paniceae, Panicoideae, Poaceae). Syst. Bot. 2007; 32(3): 531–544.

28. Aliscioni SS, Giussani LM, Zuloaga FO, Kellogg EA. A molecular phylogeny of Panicum (Poaceae: Paniceae). Test of monophyly and phylogenetic placement within the Panicoideae. Amer. J. Bot. 2003; 90(5): 796–821.

29. Kellogg EA, Hiser KM, Doust AN. Taxonomy, phylogeny, and inflorescence development of the genus Ixophorus (Panicoideae: Poaceae). Int. J. Plant Sci. 2004; 165(6): 1089–1105.

30. Hackel J, Vorontsova MS, Nanjarisoa OP, Hall RC, Razanatsoa J, Malakasi P, Besnard G. Grass diversification in Madagascar: in situ radiation of two large C_3_ shade clades and support for a Miocene to Pliocene origin of C_4_ grassy biomes. J. Biogeogr. 2018; 45: 750–761.

31. Clark LG, Zhang W, Wendel JF. A phylogeny of the grass family (Poaceae) based on ndhF sequence data. Syst. Bot. 1995; 20(4): 436–460.

32. Bess EC, Doust AN, Kellogg EA. A naked grass in the ‘‘bristle clade’’: a phylogenetic and developmental study of Panicum section Bulbosa (Paniceae: Poaceae). Int. J. Plant. Sci. 2005; 166(3): 371–381.

33. Hitchcock AS, Chase A. The North American species of Panicum. Contr. U.S. Natl. Herb. 1910; 15: 1–396.

34. Davidse G, Pohl RW. New Taxa and Nomenclatural Combinations of Mesoamerican Grasses (Poaceae). Novon. 1992; 2(2): 81–110.

35. Doyle JJ, Doyle JL. A rapid DNA isolation procedure for small quantities of fresh leaf tissue. Phytochem Bull. Bot. Soc. Amer. 1987; 19(1):11–15.

36. Olmstead RG, Sweere JA. Combining data in phylogenetic systematics: an empirical approach using three molecular data sets in the Solanaceae. Syst. Biol. 1994; 43(4): 467–481.

37. Tamura K, Stecher G, Peterson D, Filipski A., Kumar S. MEGA6: Molecular Evolutionary Genetics Analysis version 6.0. Molec. Biol. Evol. 2013; 30(12): 2725–2729.

38. Larkin MA, Blackshields G, Brown NP, Chenna R, McGettigan PA, McWilliam H, et al. Clustal W and Clustal X version 2.0. Bioinformatics. 2007; 23(21): 2947–2948.

39. Fitch WM. Toward defining the course of evolution: Minimal change for a specific tree topology. Syst. Zool. 1971; 20(4): 406–416.

40. Felsenstein J. Evolutionary trees from DNA sequences: A maximum likelihood approach. J. Molec. Evol. 1981; 17(6): 368–376.

41. Huelsenbeck JP, Crandall KA. Phylogeny estimation and hypothesis testing using maximum likelihood. Annual Rev. Ecol. Syst. 1997; 28: 437–466.

42. Huelsenbeck JP, Larget B, Miller RE, Ronquist F. Potential applications and pitfalls of Bayesian inference of phylogeny. Syst. Biol. 2002; 51(5): 673–688.

43. Goloboff PA, Farris JS, Nixon KC. TNT, a free program for phylogenetic analysis. Cladistics. 2008; 24(5): 774–786.

44. Felsenstein J. Confidence limits on phylogenies: An approach using the bootstrap. Evolution. 1985; 39(4): 783–791.

45. Stamatakis A. RAxML-VI-HPC: Maximum likelihoodbased phylogenetic analyses with thousands of taxa and mixed models. Bioinformatics. 2006; 22(21): 2688–2690.

46. Miller MA, Pfeiffer W, Schwartz T. Creating the CIPRES Science Gateway for inference of large phylogenetic trees. Proceedings of the Gateway Computing Environments Workshop (GCE), New Orleans. 2010; 01–08.

47. Stamatakis A, Hoover P, Rougemont J. A rapid bootstrap algorithm for the RAxML web-servers. Syst. Biol. 2008; 57(5): 758–771.

48. Ronquist F, Huelsenbeck JP. MrBayes 3: Bayesian phylogenetic inference under mixed models. Bioinformatics. 2003; 19(12): 1572–1574.

49. Darriba D, Taboada GL, Doallo R, Posada D. jModelTest 2: More models, new heuristics and parallel computing. Nat. Methods. 2012; 9(8): 772.

50. Rambaut A, Drummond AJ, Xie D, Baele G, Suchard MA. Posterior summarisation in Bayesian phylogenetics using Tracer 1.7. Syst. Biol. 2018; 67(5): 901–904.

51. Delfini C, Acosta JM, Pensiero JF, Aliscioni, SS. An expanded phylogeny of Setaria (Poaceae, Panicoideae, Paniceae) and its relationships within the subtribe Cenchrinae, Consejo Nacional de Investigaciones Científicas y Técnicas [dataset online]. Available from: http://hdl.handle.net/11336/163438 [accessed 1 August 2022].

52. Gallaher TJ, Peterson PM, Soreng RJ, Zuloaga FO, Li D.-Z, Clark LG, et al. Grasses through space and time: an overview of the biogeographical and macroevolutionary history of Poaceae. J. Syst. Evol. 2022; 60(3): 522–569.

53. Pilger R. Gramineae III. Unterfamilie Panicoideae. In: Engler A, Prantl K, editors, Die Natürlichen Pflanzenfamilien, vol. 14e, Leipzig: W Engelmann, Saxony; 1940. pp. 01–208.

54. Vasey G. New species of grasses. Bull. Torrey. Bot. Club. 1884; 11(6): 61–72.

55. Olorode O. Additional chromosome counts in Nigerian grasses. Brittonia. 1975; 27: 63–68.

56. Mehra PN, Sharma ML. Cytological studies in some central and eastern Himalayan grasses. II. The Paniceae. Cytologia. 1975; 1975:75–89.

57. Sarkar AK, Datta N, Mallick R, Chatterjee U. IOPB chromosome number reports LIV. Taxon. 1976; 25: 648–649.

58. Christopher J, Abraham A. Studies on the cytology and phylogeny of South Indian grasses. III. Subfamily VI: Panicoideae, tribe Paniceae. Cytologia. 1976; 41: 621–637.

59. Gadella TWJ. Chromosome number reports LVI. Taxon. 1977; 26: 257–274.

60. Dujardin M. Chromosome numbers of some tropical African grasses from western Zaire. Can. J. Bot. 1978; 56(17): 2138–2152.

61. Freitas-Sacchet AMO. Chromosome number reports LXIX. Taxon. 1980; 29: 703–704.

62. Mehra PN. Cytology of East Indian grasses. Chandigarh: PN Mehra; 1982.

63. Freitas-Sacchet AMO, Boldrini II, Born GG. Cytogenetics and evolution of the native grasses of Rio Grande do Sul, Brazil, Setaria P. Beauv. (Gramineae). Rev. Bras. Genet. 1984; 7(3): 535–548.

64. Chemisquy MA, Giussani LM, Scataglini MA, Kellogg EA, Morrone O. Phylogenetic studies favour the unification of Pennisetum, Cenchrus and Odontelytrum (Poaceae): a combined nuclear, plastid and morphological analysis, and nomenclatural combinations in Cenchrus. Ann. Bot. 2010; 106(1): 107–130.

65. Hitchcock AS. Pseudochaetochloa, a new genus of grass from Australia. J. Washington Acad. Sci. 1924; 14(21): 491–491.

66. Sauer JD. Revision of Stenotaphrum (Gramineae: Paniceae) with attention to its historical geography. Brittonia. 1972; 24(2): 202–222.

67. Webster RD. The Australian Paniceae (Poaceae). Berlin: J. Cramer; 1987.

68. Pilger R. Acritochaete, eine neue Gramineen-Gattung aus Afrika. Bot. Jahrb. Syst. 1902; 32(1): 53–55.

69. Cotton JL, Wysocki WP, Clark LG, Kelchner SA, Pires JC, Edger PP, et al. Resolving deep relationships of PACMAD grasses: A phylogenomic approach. BMC Plant Biology. 2015; 15: 01–11.

70. Burke SV, Wysocki WP, Zuloaga FO, Craine JM, Pires JC, Edger PP, et al. Evolutionary relationships in Panicoid grasses based on plastome phylogenomics (Panicoideae; Poaceae). BMC Plant Biology. 2016; 16: 01–11.

71. Saarela JM, Burke SV, Wysocki WP, Barrett MD, Clark LG, Craine JM, et al. A 250 plastome phylogeny of the grass family (Poaceae): Topological support under different data partitions. PeerJ. 2018; 2018: e4299.

72. Acosta JM, Zuloaga FO, Reinheimer R. Nuclear phylogeny and hypothesized allopolyploidization events in the subtribe Otachyriinae (Paspaleae, Poaceae). Syst. Biodivers. 2019; 17(3): 277–294.

73. Huang W, Zhang L, Columbus JT, Hu Y, Zhao Y, Tang L, et al. A well-supported nuclear phylogeny of Poaceae and implications for the evolution of C_4_ photosynthesis. Mol. Plant. 2022; 15(4): 755–777.

74. Estep MC, McKain MR, Diaz DV, Zhong J, Hodge JG, Hodkinson, TR, et al. Allopolyploidy, diversification, and the Miocene grassland expansion. Proc. Natl. Acad. Sci. U.S.A. 2014; 111(42): 15149–15154.

75. Kellogg EA. Has the connection between polyploidy and diversification actually been tested? Curr. Opin. Pl. Biol. 2016; 30: 25–32.

76. Sang T. Utility of low-copy nuclear gene sequences in plant phylogenetics. Crit. Rev. Biochem. Mol. Biol. 2002; 37(3): 121–147.

